# Innate, translation-dependent silencing of an invasive transposon in Arabidopsis

**DOI:** 10.1101/2021.06.29.450179

**Authors:** Stefan Oberlin, Rajendran Rajeswaran, Marieke Trasser, Verónica Barragán-Borrero, Michael A. Schon, Alexandra Plotnikova, Lukas Loncsek, Michael D. Nodine, Arturo Marí-Ordóñez, Olivier Voinnet

## Abstract

Co-evolution between hosts’ and parasites’ genomes shapes diverse pathways of acquired immunity based on silencing small (s)RNAs. In plants, sRNAs cause heterochromatinization, sequence-degeneration and, ultimately, loss-of-autonomy of most transposable elements (TEs). Recognition of newly-invasive plant TEs, by contrast, involves an innate antiviral-like silencing response. To investigate this response’s activation, we studied the single-copy element *EVADÉ* (*EVD*), one of few representatives of the large *Ty1/Copia* family able to proliferate in Arabidopsis when epigenetically-reactivated. In *Ty1/Copia-*elements, a short subgenomic mRNA (*shGAG*) provides the necessary excess of structural GAG protein over the catalytic components encoded by the full-length genomic *flGAG-POL*. We show here that the predominant cytosolic distribution of *shGAG* strongly favors its translation over mostly-nuclear *flGAG-POL*, during which an unusually intense ribosomal stalling event coincides precisely with the starting-point of sRNA production exclusively on *shGAG.* mRNA breakage occurring at this starting-point yields unconventional 5’OH RNA fragments that evade RNA-quality-control and concomitantly likely stimulate RNA-DEPENDENT-RNA-POLYMERASE-6 (RDR6) to initiate sRNA production. This *hitherto*-unrecognized “translation-dependent silencing” (TdS) is independent of codon-usage or GC-content and is not observed on TE remnants populating the Arabidopsis genome, consistent with their poor association, if any, with polysomes. We propose that TdS forms a primal defense against *de novo* invasive TEs that underlies their associated sRNA patterns.

## Introduction

Transposable elements (TEs) colonize and threaten the integrity of virtually all genomes (Huang *et al*, 2012). Chromosomal rearrangements caused by their highly-repetitive nature (Fedoroff, 2012) are usually circumvented by cytosine methylation and/or histone-tail modifications at their loci-of-origin. The ensuing heterochromatic DNA is not conducive to transcription by RNA Pol II, bringing TEs into an epigenetically silent transcriptional state (Allshire & Madhani, 2018). This “transcriptional gene silencing” (TGS) is observed at the majority of TE loci in plants, including the model species *Arabidopsis thaliana,* and causes, over evolutionary times, accumulating mutations resulting in mostly degenerated, non-autonomous entities (Vitte & Bennetzen, 2006; Civáň *et al*, 2011). Nonetheless, the genome-invasiveness of these remnants remains evident by their methyl cytosine-marked DNA, which is perpetuated over generations by METHYL-TRANSFERASE 1 (MET1), among other factors. MET1 reproduces symmetrical methylation sites from mother- to daughter-strands during DNA replication (Kankel *et al*, 2003) aided by the (hetero)chromatin remodeler DEFICIENT IN DNA METHYLATION 1 (DDM1) (Saze *et al*, 2003; Zemach *et al*, 2013).

Loss of MET1 or DDM1 functions in *Arabidopsis* leads to genome-wide demethylation, transcriptional reactivation of many TE remnants, and mobilization of a small portion of intact, autonomous TEs (Mirouze *et al*, 2009; Tsukahara *et al*, 2010). Their proliferation together with genome-wide deposition of aberrant epigenetic marks likely explains why *met1* and *ddm1* mutants accumulate increasingly severe genetic and phenotypic burdens over inbred generations (Vongs *et al*, 1993). However, such secondary events can be avoided by backcrossing the first homozygous generation of *ddm1-* or *met1-*derived mutants with wild type plants, upon which continuous selfing of F2 plants creates “epigenetic recombinant inbred lines” (epiRILs). These harbor only mosaics of de-methylated DNA while maintaining wild-type (WT) MET1 and DDM1 functions (Teixeira *et al*, 2009; Reinders *et al*, 2009a). One such *met1* epiRIL, epi15, endows epigenetic reactivation of the autonomous, long terminal repeat (LTR) retroelement *EVADÉ* (*EVD*) in the*Ty1/Copia* family, which is one of the most proliferative families in plants (Vitte & Panaud, 2005). Of the two *EVD* copies in the *Arabidopsis* Col-0 genome, only one is reactivated in epi15 (Marí Ordóñez *et al*, 2013). By providing a proxy for a *de novo* genomic invasion, this reactivation granted a unique opportunity to grasp how, over multiple inbred generations, newly invasive TEs might be detected and eventually epigenetically silenced (Marí Ordóñez *et al*, 2013).

We found that *EVD* is initially confronted to post-transcriptional gene silencing (PTGS) akin to that mounted against plant viruses (Voinnet, 2005; Marí Ordóñez *et al*, 2013). Antiviral RNA-DEPENDENT RNA POLYMERASE 6 (RDR6) produces cytosolic, long double-stranded (ds)RNAs from *EVD*-derived transcripts, which are then processed by DCL4 or DCL2, two of the four *Arabidopsis* Dicer-like RNase-III enzymes, into populations of respectively 21- and 22-nt small interfering (si)RNAs. However, despite their loading into the antiviral PTGS effectors ARGONAUTE1 and ARGONAUTE2 (AGO1/2), they do not suppress expression of *EVD*’s increasingly more abundant genomic copies. This ultimately gives way to DCL3, instead of DCL4/2, to process the RDR6-made long dsRNAs into 24-nt siRNAs. In association with AGO4-clade AGOs, these species guide RNA-directed DNA methylation (RdDM) of *EVD* copies. Initially localized within the *EVD* gene body, it later spreads into the LTRs to eventually shut down the expression of *EVD* genome-wide via TGS (Marí Ordóñez *et al*, 2013).

A key, unsolved question prompted by this proposed suite of events pertains to the mechanisms whereby RDR6 is initially recruited onto *EVD,* and more generally on newly invasive TEs, during the primary antiviral-like silencing phase. “Homology-” or “identity”-based silencing entails sequence complementarity between TE transcripts and host-derived small RNAs. Loaded into AGOs, they likely attract RDR6 concomitantly to silencing execution. One such type of PTGS occurs with TEs reactivated in *ddm1*/*met1* mutants, which, by displaying complementarity mostly to host-encoded microRNAs, spawn “epigenetically-activated siRNAs” (easiRNAs) in an AGO1-dependent manner (Creasey *et al*, 2014). easiRNA production likely entails substantial co-evolution between host and TE genomes (Sarazin & Voinnet, 2014) because miRNAs usually target short and highly conserved TE regions, including the primer-binding sites required for retroelements’ reverse-transcription (RT) (Šurbanovski *et al*, 2016; Borges *et al*, 2018). Another form of acquired immunity underlying identity-based silencing is conferred by siRNAs derived from relics of previous genome invasions by the same or sequence-related TE(s) (Fultz & Slotkin, 2017).

New intruder TEs are unlikely to engage either form of identity-based silencing, as indeed noted for *EVD* (Creasey *et al*, 2014). Thus, RDR6-dependent PTGS initiation should involve intrinsic features of the TEs themselves (Sarazin & Voinnet, 2014). In the yeast *Cryptococcus neoformans*, stalled spliceosomes on suboptimal TE introns provide an opportunity for an RDR-containing complex to co-transcriptionally initiate such innate PTGS (Dumesic *et al*, 2013). Studies of transgene silencing in plants (Luo & Chen, 2007; Thran *et al*, 2012) have advocated other possible mechanisms, though none has yet been linked to epigenetically reactivated TEs. These studies describe how uncapped, prematurely terminated or non-polyadenylated transcripts might stimulate RDR activities when they evade or overwhelm RNA quality control (RQC) pathways that normally degrade these “aberrant” RNAs (Parent *et al*, 2015; Gy *et al*, 2007b; Herr *et al*, 2006). A recent model also contends that widespread translation-coupled RNA degradation as a consequence of suboptimal codon usage and low GC content might trigger RDR-dependent silencing in plants (Kim *et al*, 2021). Initiation of innate PTGS in the context of *EVD* likely ties in with an unusual process of splicing-coupled premature cleavage and poly-adenylation (PCPA) shared by *Ty1/Copia* retroelements to optimize protein expression from their compact genomes (Oberlin *et al*, 2017).On the one hand, an unspliced and full-length (*fl*) *GAG-POL* isoform codes for a polyprotein processed into protease, integrase/reverse-transcriptase-RNase and GAG nucleocapsid components. On the other hand, a spliced and prematurely terminated short *(sh) GAG* subgenomic isoform is solely dedicated to GAG production. Though less abundant than the *flGAG-POL* mRNA, *shGAG* is substantially more translated (Oberlin *et al*, 2017). This presumably results in a molar excess of structural GAG for viral-like particle (VLP) formation compared to Pr-IN-RT-RNase required for reverse-transcription (RT) and, ultimately, mobilization (Oberlin *et al*, 2017; Lee *et al*, 2020). Supporting the notion that genome expression of *Ty1/Copia* elements influences PTGS initiation, *EVD*-derived RDR6-dependent *s*iRNAs do not map onto the unspliced *flGAG-POL* mRNA, but instead specifically onto the spliced *shGAG* transcript of which, intriguingly, they only cover approximately the 3’ half (Oberlin *et al*, 2017).

Here, we show that differential subcellular distribution of the two mRNA isoforms due to splicing-coupled PCPA accounts for the peculiar *EVD* siRNA distribution and activity patterns. While the *flGAG-POL* isoform remains largely nuclear, the *shGAG* mRNA is enriched in the cytosol and endows vastly disproportionate translation of *shGAG* over *flGAG-POL*. However, a previously uncharacterized innate PTGS process accompanies active *shGAG* translation, manifested as a discrete and unusually intense ribosome stalling event independent of codon usage or GC content, among other tested parameters. Ribosome stalling coincides precisely with the starting point of *shGAG* siRNA production and maps to the 5’ ends of discrete, *shGAG*-derived RNA breakage fragments. These harbor unconventional 5’OH *termini* that prevent their RQC-based degradation via 5’P-dependent XRN4 action (Stevens, 2001; Peach *et al*, 2015). Based on the well-documented substrate competition between XRN4 and RDR6(Gazzani, 2004; Gy *et al*, 2007b; Gregory *et al*, 2008; Moreno *et al*, 2013; Martínez de Alba *et al*, 2015), we suggest that the 5’OH status of breakage fragments licenses their conversion into dsRNA by RDR6, thereby initiating PTGS of *EVD*. We further show that splicing-coupled PCPA suffices to recapitulate this “translation-dependent silencing” (TdS) in reporter-gene settings. Given that *Ty1/Copia* retroelements share a PCPA-based genome expression strategy (Oberlin *et al*, 2017), we contend that TdS forms a generic and primal defense against *de novo* invasive TEs that shapes the siRNA patterns initially associated with them.

## Results

### *shGAG* is the main source and target of *EVD*-derived siRNAs

*Arabidopsis* lines constitutively overexpressing an LTR-deficient but otherwise intact form of *EVD* driven by the 35S promoter (*35S:EVD_wt_*) recapitulate the restriction of *EVD* siRNA to the 3’ part of the *shGAG* sequence(Marí Ordóñez *et al*, 2013; Oberlin *et al*, 2017) (Fig.1A-B). We explored *EVD* transcripts levels in *35S:EVD_wt_* in WT (siRNA-proficient) as opposed to *rdr6* (siRNA-deficient) background (Fig.1B, S1A). Both in RNA blot and qRT-PCR analyses, the spliced *shGAG* mRNA levels were increased in *rdr6* compared to WT, whereas those of unspliced *flGAG-POL* were globally unchanged (Fig.1C-D). Accordingly, accumulation of the GAG protein – mainly produced *via shGAG* translation (Oberlin *et al*, 2017) – was higher in *rdr6* compared to WT background (Fig.1E). Essentially identical results were obtained upon epigenetic reactivation of endogenous *EVD* in non-transgenic Arabidopsis with the *ddm1* single-*versus ddm1 rdr6* double mutant background (Fig.S1B-E). Following *EVD* mobilization from an early (F8) to a more advanced (F11) epi15 inbred generation (Marí Ordóñez *et al*, 2013) revealed that its progressively increased copy number correlates with progressively higher steady-state levels of *EVD*-derived transcripts and *EVD-*derived siRNAs (Fig.S1F-G). Again, these siRNAs disproportionately target the *shGAG* relative to *flGAG-POL* mRNA from F8 to F11 (Fig.S1H). Collectively, these results indicate that PTGS activated *de novo* by *EVD* is both triggered by, and targeted against, the spliced *shGAG* mRNA. Therefore, features associated with *shGAG,* but not *flGAG-POL,* likely stimulate RDR6 recruitment, which we explored by testing current models for PTGS initiation from TEs and transgenes.

**Figure 1.**
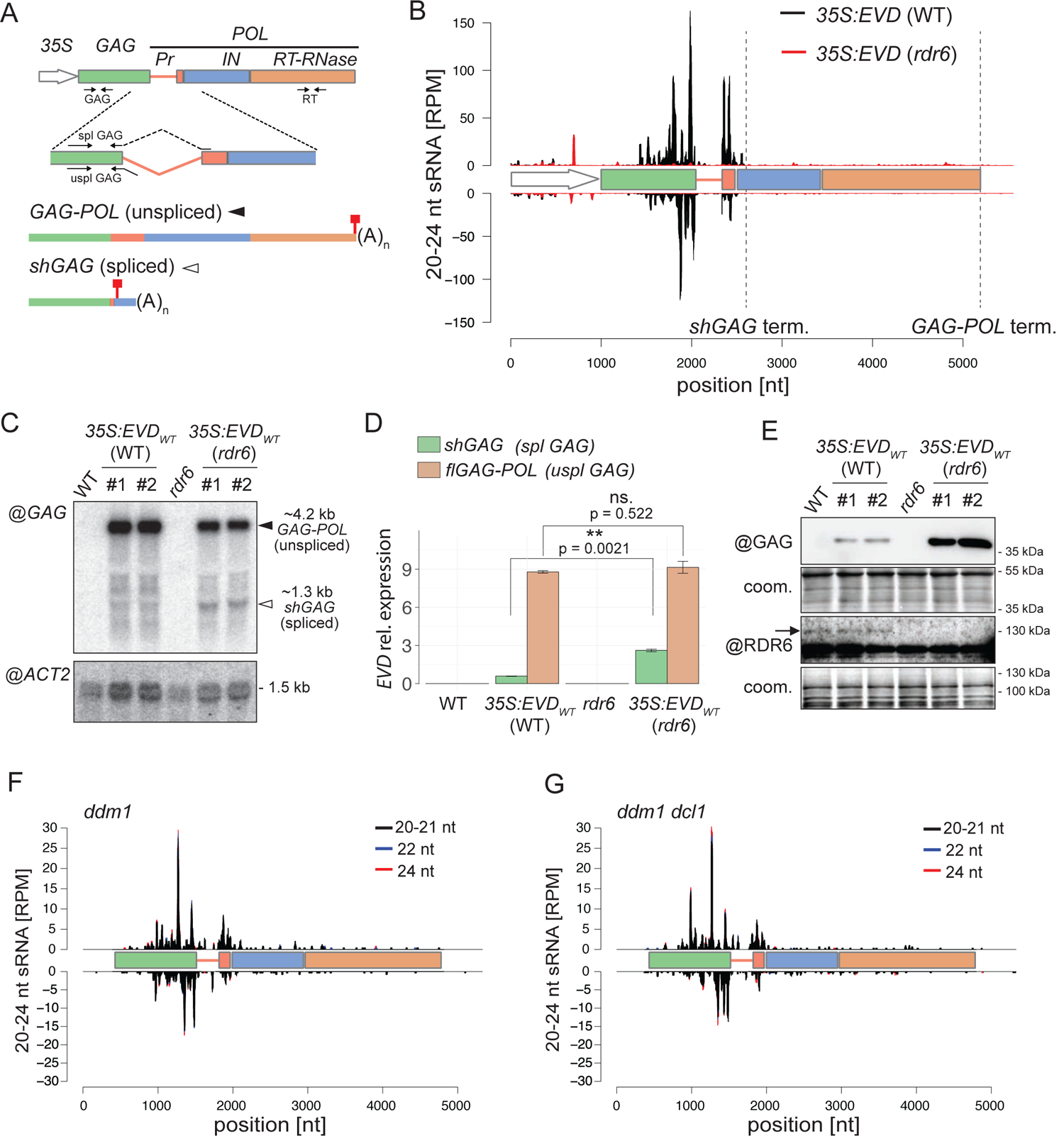
*EVD shGAG* is both a trigger and a target of RDR6-dependent but miRNA-independent siRNAs. **(A)** *EVD flGAG-POL* and spliced *shGAG* mRNAs are distinguishable using specific PCR primer sets (arrows) for quantification and northern analysis. (*35S*) *Cauliflower Mosaic Virus 35S promoter*, (*Pr*) protease, (*IN*) integrase, (*RT-RNase*) reverse-transcriptase-RNase; red squares: stop codons. **(B)** sRNA-seq reads profile of *EVD* expressed from *35S:EVD_WT_* in *WT* (black) or *rdr6* (red). (RPM) reads per million. Positions are indicated in nucleotides (nt) from the start of the *35S* sequence. Dashed vertical lines: *shGAG* and *GAG-POL* 3’ ends. (C) Northern analysis of *EVD* RNA isoforms using a probe for the *GAG* region or for *ACTIN2* (*ACT2*) as a loading control. **(D)** qPCR quantification of *shGAG* and *flGAG-POL* normalized to *ACT2* and to *GLYCERALDEHYDE-3-PHOSPHATE DEHYDROGENASE C SUBUNIT* (*GAPC*) levels. qPCR was performed on n=3 biological replicates; bars: standard error. (**) = p-value < 0.01 (two-sided t-test between indicated values). **(E)** Western analysis of GAG and RDR6 with Coomassie (coom.) staining as a loading control. **(F-G)** sRNA-seq profiles from *EVD* de-repressed in the *ddm1* (F) or *ddm1 dcl1* (G) backgrounds. Different siRNA size categories are stacked. Nomenclature as in (B).

### *shGAG* siRNA production is likely miRNA-independent

Though unlikely (Creasey *et al*, 2014; Sarazin & Voinnet, 2014), we first considered that production of RDR6-dependent siRNAs from *shGAG* might require its cleavage by miRNAs via the easiRNA pathway (Creasey *et al*, 2014). *Arabidopsis* miRNA biogenesis depends on DCL1 and the dsRNA-binding protein HYL1, among other factors (Brodersen & Voinnet, 2006). Analyses of publicly available sRNA-seq data (Creasey *et al*, 2014) showed, however, that epigenetically reactivated *EVD* spawns qualitatively and quantitatively identical *shGAG-*only siRNAs in both *ddm1* single and *ddm1 dcl1* double mutants (Fig.1F-G). Moreover, levels of *shGAG* siRNA, *shGAG* mRNA, and GAG protein remained unchanged in *35S:EVD_wt_* plants with either the WT, hypomorphic *dcl1-11* or loss-of-function *hyl1-2* background (Fig.S2A-D). By contrast and as expected, production of trans-acting (ta)siRNAs, which is both miRNA- and RDR6-dependent, was dramatically reduced and the levels of tasiRNA precursors and target transcripts enhanced in both mutant backgrounds (Fig.S2). Therefore, RDR6 recruitment to the spliced *shGAG* mRNA is unlikely to involve endogenous miRNAs *via* an identity-based mechanism. We then explored known innate processes of PTGS initiation instead.

### Splicing-coupled premature cleavage and poly-adenylation suffices to spawn

#### *EVD*-like siRNA accumulation and activity patterns

Some cases of transgene-induced PTGS correlate with a lack of polyadenylation due to aberrant RNA transcription (Luo and Chen 2007). We ruled that this feature underlies *EVD*-derived siRNA production because *shGAG* displays no overt polyadenylation defects regardless of the onset of PTGS (Fig.S3). Next, we considered splicing defects, such as inaccurate splicing or spliceosome stalling, and premature transcriptional termination as possible PTGS triggers, two processes previously independently linked to innate, RDR-dependent siRNA production in plants and fungi ((Dalakouras *et al*, 2019; Dumesic *et al*, 2013). *Ty1/Copia* elements have introns that are significantly longer than those of Arabidopsis genes. Moreover, *shGAG* undergoes atypical splicing-coupled PCPA (Oberlin *et al*, 2017). When engineered between the *GFP* and *GUS* sequences of a translational fusion, the *shGAG* intron and proximal PCPA signal spawn unspliced *flGFP-GUS* and spliced *GFP-*only (*shGFP*) mRNAs in the *Arabidopsis* line *35S:GFP*-*EVD_int/ter_*-*GUS* (Oberlin *et al*, 2017) (Fig.2A-B;Fig.S4A). Since this artificial system recapitulates the production of respectively *flGAG-POL* and *shGAG*, we asked if an *EVD*-like siRNA pattern was likewise reproduced.

**Figure 2.**
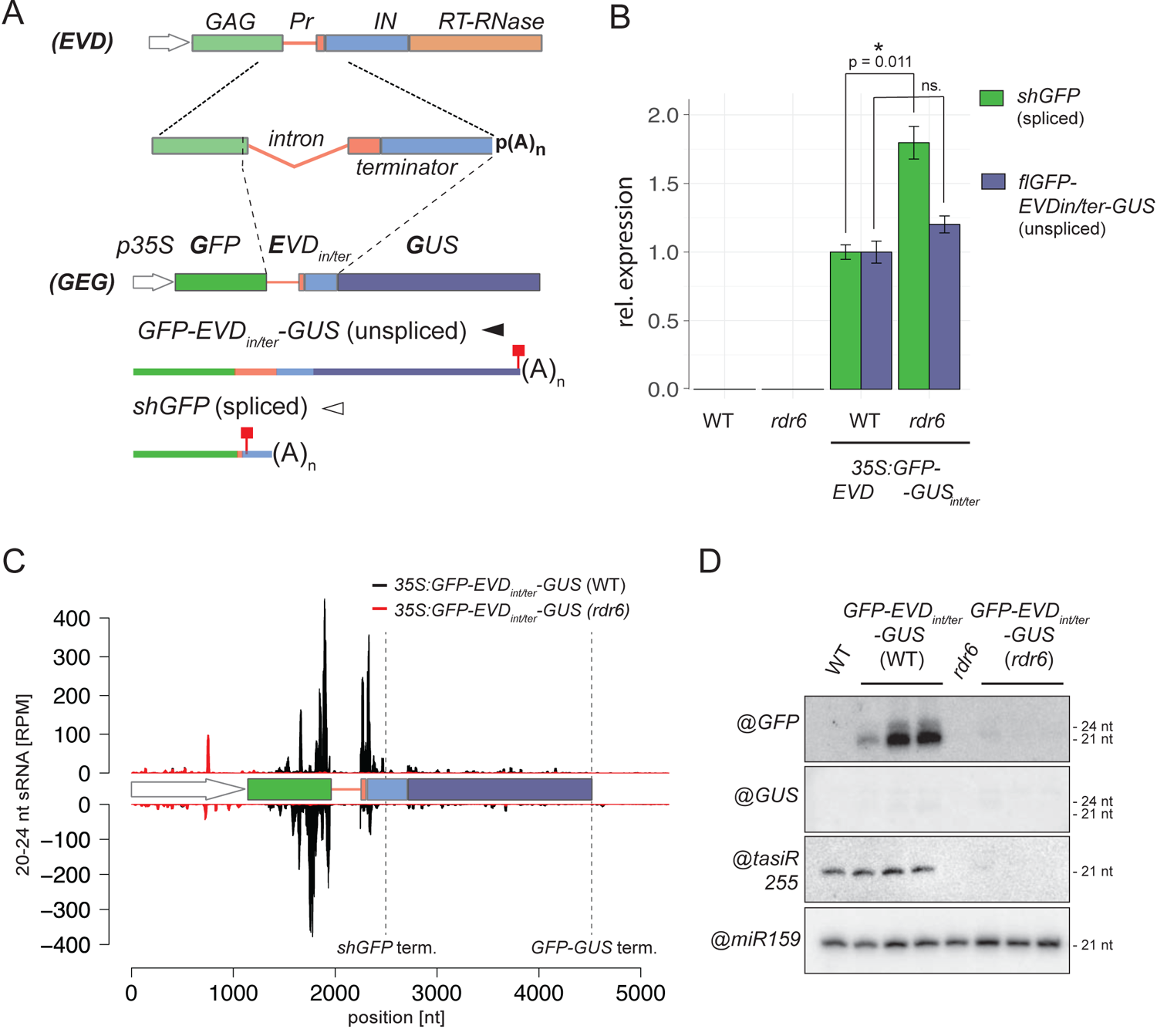
The *EVD* intron and terminator suffice to initiate PTGS. **(A)** The *35S:GFP-EVDint/ter-GUS* fusion was made by introducing the *EVD* intron and proximal *shGAG* terminator between the *GFP* and *GUS* coding sequence. Like *EVD*, it spawns full length unspliced and short spliced mRNAs. Red squares: stop codons. **(B)** Expression levels of *shGFP* (spliced) and *GFP-EVDint/ter-GUS* (unspliced) transcripts, relative to *ACT2* and AT4G26410 (*RHIP1*), in the WT or *rdr6* background. qPCR was performed on three biological replicates and error bars represent the standard error on. (*) = p-value < 0.05 (two-sided t-test against corresponding controls). **(C)** sRNA-seq profile mapped on the genomic *35S:GFP-EVDint/ter-GUS locus.* (RPM) Reads per million. Positions indicated in nucleotides (nt) from the start of the 35S sequence. Dashed vertical lines: *shGFP* and *GFP-GUS* 3’ ends. **(D)** Low molecular weight RNA analysis of the *GFP*- and *GUS*-spanning regions. tasiRNA255 is a control for the rdr6 mutation and miR159 provides a loading control.

The majority of RDR6-dependent 21-nt siRNAs mapped to the *GFP*, but not the *GUS* region downstream of the PCPA signal (Fig.2C-D) suggesting that, just like *shGAG* in *EVD*, the spliced *shGFP* mRNA is the main source of siRNAs in *GFP*-*EVD_int/ter_*-*GUS*. Accordingly, and similar to *EVD* (Oberlin *et al*, 2017), many siRNAs spanned the exon-exon junction of *GFP*-*EVD_int/ter_*-*GUS* (Fig.S4B). Moreover, *shGFP,* unlike *flGFP-GUS,* over-accumulated in *GFP*-*EVD_int/ter_*-*GUS* plants with the *rdr6* background (Fig.2B, Fig.S4A), indicating that only *shGFP* is efficiently targeted by PTGS (Fig.2B). Therefore, in the reconstituted setting, the intron and PCPA signal found in *shGAG* suffice to spawn RDR6-dependent siRNAs displaying accumulation and activity patterns resembling those generated in the authentic *EVD* context (Fig.1A-D).

### Neither splicing nor intron-retention *per se* initiate RDR6 recruitment

The above result prompted us to investigate a potential facilitating role for splicing in *shGAG* siRNA biogenesis or, conversely, a role for intron-retention in inhibiting RDR6 recruitment to *flGAG-POL*. We used previously engineered Arabidopsis *EVD*-overexpression lines with a point-mutated U1 snRNP-binding site (35S:*EVD_mU1_*) or a fully deleted intron (35S:*EVD_Δi_*) (Oberlin *et al*, 2017) (Fig.3A-C). *35S:EVD_Δi_* spawns fully matured s*hGAG* transcripts that do not associate with the spliceosome, leading exclusively to prematurely terminated and polyadenylated mRNA species with a stop codon (Oberlin *et al*, 2017) (Fig.3B, 3D, 3E, Fig.S5A). However, lack of the intron, and hence splicing, did not prevent RDR6-dependent siRNA production from *35S:EVD_Δi_*, which was comparable to that of *35S:EVD_wt_* (Fig.3E). Moreover, the *shGAG* mRNA and GAG protein levels from *35S:EVD_Δi_* were higher in an *rdr6* compared to WT background (Fig.3D-F), indicating that *EVD’s* unconventional splicing is unlikely to underpin *shGAG* siRNA production.

**Figure 3.**
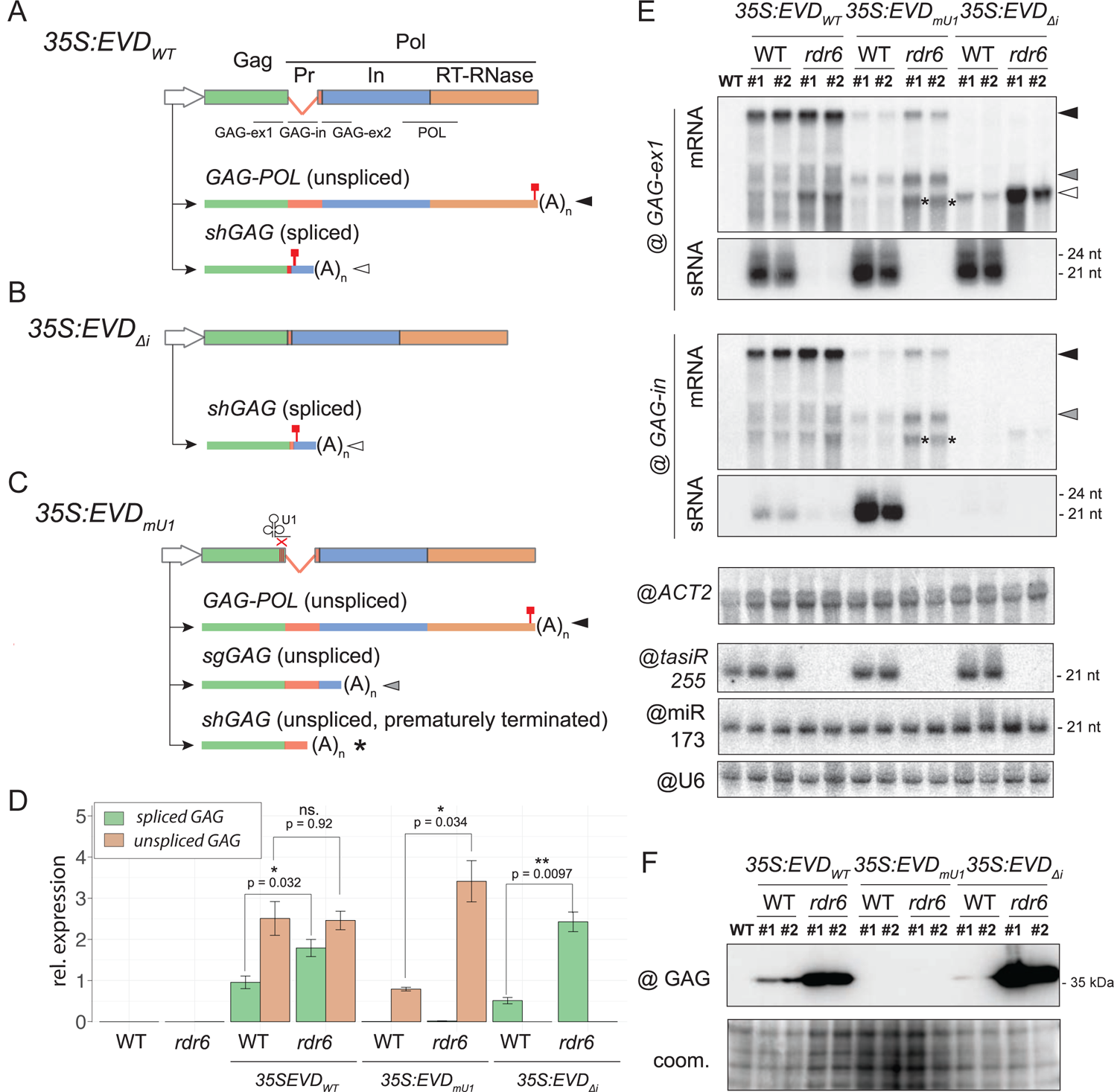
Impact of splicing and premature termination on *EVD* silencing. **(A-C)** Constructs and isoforms transcribed from *35S:EVDwt* (A), *35S:EVDL1intron* (B) and 35S:EVDmU1 (C). Probes for northern analysis of *GAG* exon 1 (GAG-ex1), intron (GAG-in), exon2 (GAG-ex2) and the POL region are depicted with black lines. **(D)** Relative expression levels of spliced and unspliced transcripts in the three *EVD* constructs relative to *ACT2*. qPCR was performed on three biological replicates and error bars represent the standard error. (ns.) = non-significant, (*) = p-value < 0.05, (**) = p-value < 0.01, (two-sided t-test between indicated samples/targets). **(E)** High and low molecular-weight RNA analysis of *EVD GAG* (GAG-ex1) and *EVD* intron (GAG-in) in two independent T1 bulks from each indicated line. The filled arrows on the right hand-side or with an asterisk on the blots correspond to the transcripts depicted in (A-C). *ACT2*: loading control for mRNAs; tasiR255, miR173 and U6: loading controls for sRNAs. Hybridizations for GAG-ex2 and POL probes are found in Supp. Fig.5A. **(F)** Western analysis of the GAG protein with Coomassie (coom.) staining as a loading control.

To test the alternative possibility that intron-retention or specific sequences within the *EVD* intron prevent siRNA biogenesis from *flGAG-POL*, we analyzed the siRNAs from 35S:*EVD_mU1_*. Impeding U1 binding and its inhibitory action on PCPA causes a complete lack of splicing in *EVD_mU1_* (Fig.3C-D). This generates short unspliced transcripts, alternatively terminated at the cognate *shGAG* terminator or at an intronic cryptic site previously mapped by 3’ RACE (Oberlin *et al*, 2017), both detected here by northern analysis (Fig.3C-E, Fig.S5A). Both alternatively terminated transcripts likely undergo translation, albeit largely unproductively (Fig.3F), because low levels of cryptic GAG translation products were detectable in *rdr6* compared to WT (Fig.S5B). *EVD_mU1_* bestowed RDR6-dependent siRNA production expanding – as expected from its non-spliceable nature – into the retained intron sequence (Fig.3E). The near-complete lack of siRNAs downstream of the intron (Fig.S5A), by contrast, suggested that both cryptically terminated *shGAG* transcripts are mainly involved in recruiting RDR6. Therefore, even though the *shGAG* intron and PCPA signal suffice to trigger PTGS from *EVD* and *GFP-EVD_int/ter_*-*GUS* (Figs.1-2), neither splicing nor intron-retention *per se* seem to initiate PTGS. This suggests that splicing-coupled PCPA does not co-transcriptionally condition the sensitivity of *shGAG* to RDR6 but, rather, downstream in the gene expression pathway.

### RDR6 recruitment onto *shGAG* likely requires translation

Splicing-coupled PCPA, conserved among Arabidopsis *Ty1/Copia* elements, correlates with the over-representation of *shGAG* on polysomes as opposed to the paradoxically more abundant *flGAG-POL* (Oberlin *et al*, 2017). However, among the *ddm1-* or *met1-* reactivated *Ty1/Copia* elements sharing the same genome expression strategy, only *EVD* spawns detectable RDR6-dependent *shGAG* siRNAs (Oberlin *et al*, 2017), prompting us to explore the basis for this difference. Polysome association, independently of translation efficiency, is the most decisive prerequisite for any given RNA to engage the translation machinery. For instance, many non-coding RNAs are mostly nuclear (Khanduja *et al*, 2016), and aberrant (*e.g.* uncapped and/or poly(A)^-^) mRNAs are actively degraded by RQC, both of which explain their general absence from polysomes (Doma & Parker, 2007). We conducted genome-wide correlation analyses between steady-state transcript accumulation, polysome association, and siRNA levels of reactivated TEs in the *ddm1* versus *ddm1 rdr6* background by calculating the ratio of polysome-associated *versus* total mRNA levels. The same approach was applied to Arabidopsis protein-coding compared to non-coding RNAs used as references (Oberlin *et al*, 2017). This analysis revealed two distinct TE populations according to the levels of associated RDR6-dependent siRNAs. On the one hand, approximately ¾ of *ddm1* de-repressed TEs (530/674) display varying degrees of polysome association, some within the range of protein-coding genes (Fig. 4A, quartiles 1-3). However, RDR6-dependent siRNA production does not accompany their reactivation presumably because of their low expression levels (Fig.4B, quartiles 1-3). The remaining ¼ (144/674) of TEs spawn RDR6-dependent siRNAs, correlating with higher RNA expression levels (Fig.4A-B quartile 4). Nonetheless, unlike those of quartiles 1-3, these TEs, almost exclusively composed of degenerated *LTR/Gypsy* elements (*i.e.* elements shorter than-full-length reference ORFs; Fig.4C), resemble non-coding RNAs in being poorly associated with polysomes, if at all (Fig. 4A, quartile 4). By contrast, *EVD* is the sole *LTR/Copia* element within quartile 4, in which it is one of the most strongly polysome-associated elements that concurrently spawn 21-22-nt siRNAs. Furthermore, when the two *EVD* isoforms are considered separately, *shGAG* emerges as a clear outlier by being associated with polysomes to the same extent as protein-coding mRNAs, (Fig.4A, quartile 4, inlay). *flGAG-POL,* by contrast, displays low polysome association albeit higher than most degenerated *LTR/Gypsy* elements populating quartile 4. In summary, *shGAG,* compared to *flGAG-POL,* is both vastly overrepresented on polysomes (Oberlin *et al*, 2017) and is the major, if not unique source of *EVD*-derived siRNAs (Fig.1, S1, Fig.3). This analysis suggests, therefore, that translation is the step stimulated by splicing-coupled PCPA of *shGA*G, upon which RDR6 is recruited specifically onto this mRNA isoform.

**Figure 4.**
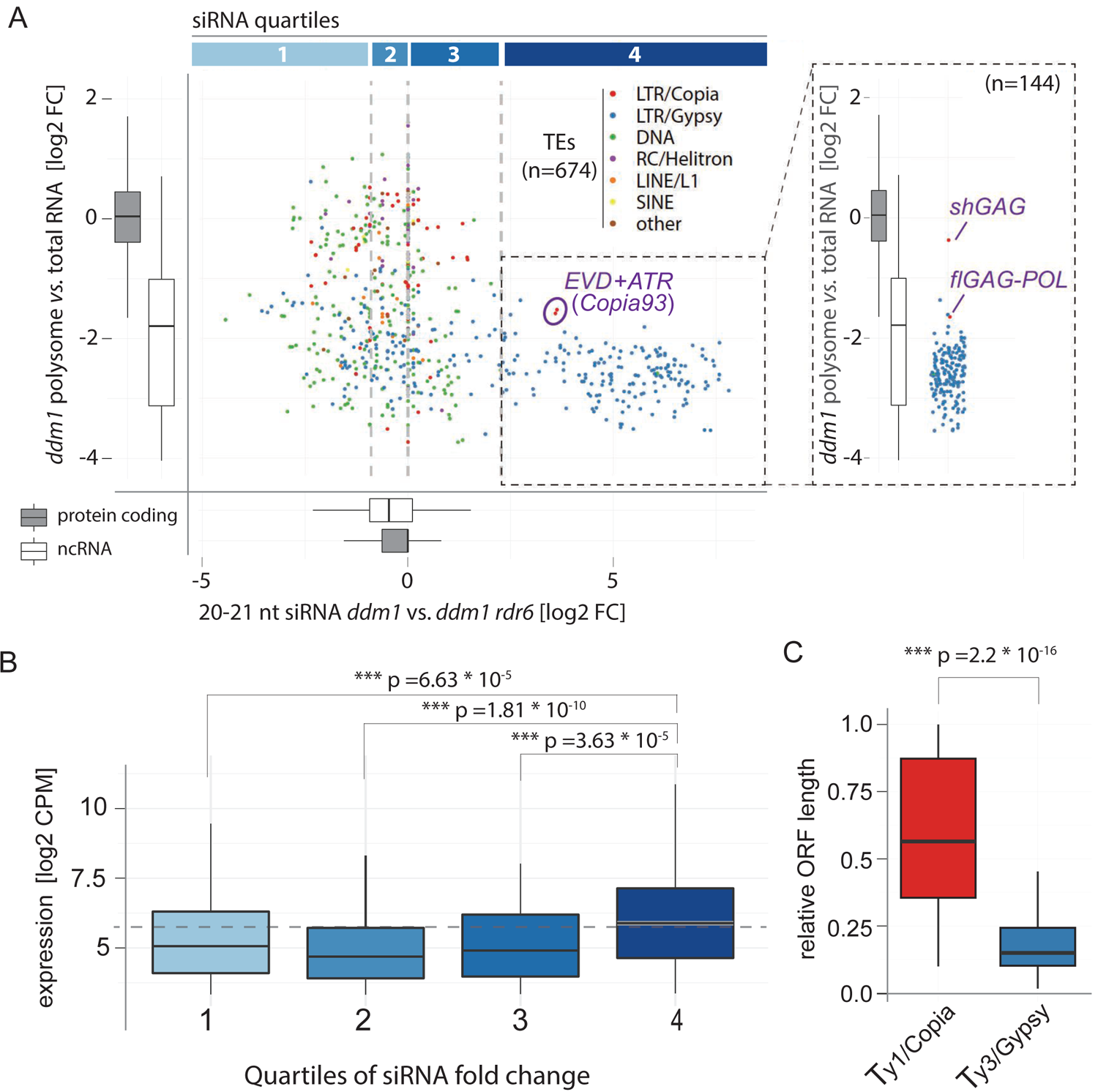
Expression, but not translation, is associated with RDR6 activity on most *ddm1*-reactivated TEs except *EVD*. polysome libraries vs. total RNA) and RDR6-dependent siRNA levels of TEs found de-repressed in *ddm1* (brief description polysome association and RDR6-dependent siRNA levels of protein coding and non-coding transcripts are displayed as boxplots. *Copia93* elements: *EVD* (AT5G17125) + *ATR* (AT1G34967), are circled. Inlet: Polysome association score of TEs in quartile 4, *EVD* mRNA isoforms are displayed separately. **(B)** Boxplots of RNA expression levels of TEs in *ddm1* from the quartiles in (A). In all panels: (***) = p-value < 0.001, (Wilcoxon rank-sum test against labelled controls or protein coding gene cohort). **(C)** ORF length of *Ty1/Copia* and *Ty3/Gypsy* elements expressed in *ddm1* relative to their genomic length.

### Splicing-coupled PCPA promotes selective translation and PTGS initiation from *shGAG*-like mRNA isoforms

To test if differential translation due to splicing-coupled PCPA indeed underlies siRNA production from *shGAG* as opposed to *flGAG-POL*, we used *GFP*-*EVD_int/ter_*-*GUS,* from which the two *EVD* RNA isoforms and associated siRNA production/activity patterns are recapitulated (Fig.2). Of the *shGAG*-like *shGFP-* and *flGAG-POL*-like *flGFP-GUS-* mRNAs, only the former produced a detectable protein under the form of free GFP (Fig.S4C-D) despite accumulation of both mRNAs (Fig.2A, Fig.S4A). Free GFP levels were increased in the *rdr6* background (Fig.S4C), coinciding with increased *shGFP-* but unchanged *flGFP-GUS*-mRNA levels (Fig.2B). The lack of detectable GFP-GUS fusion protein – the expected product of *flGFP-GUS*– in either WT or *rdr6* backgrounds (Fig.2D, Fig.S4C-D) was not due to intrinsically poor translatability. Indeed, GFP-GUS was the sole protein detected in independent lines undergoing RDR6-dependent PTGS of *35S:GFP-GUS*, a construct identical to *35S:GFP*-*EVD_int/ter_*-*GUS,* save the *shGAG* intron and PCPA signal (Fig.5A-B). As expected, the GFP-GUS fusion protein and *GFP-GUS* mRNA levels were strongly enhanced in the *rdr6* versus WT background (Fig.5A-B). Yet, in contrast to *GFP*-*EVD_int/ter_*-*GUS*, from which siRNAs are restricted to *shGFP,* the siRNAs from *GFP-GUS* encompassed both the *GFP* and *GUS* sequences (Fig.5A). These results therefore indicate that splicing-coupled PCPA promotes selective translation of, and PTGS initiation from, *shGAG*-like as opposed to *flGAG-POL*-like mRNA isoforms.

**Figure 5.**
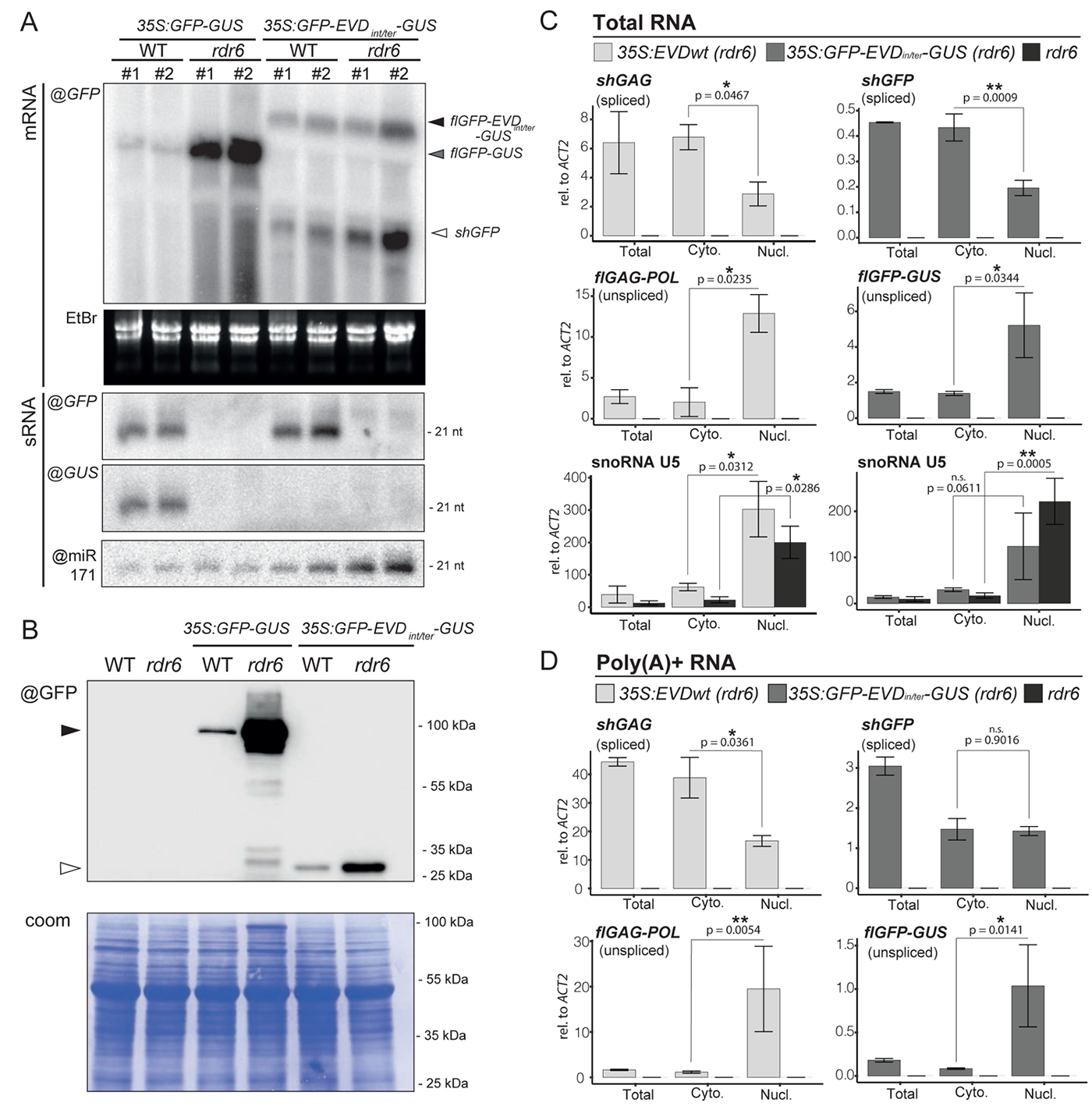
Splicing promotes translation and siRNA biogenesis from short-spliced mRNAs by influencing nucleocytoplasmic distribution of RNA isoforms. **(A)** Comparison of RNA isoforms and sRNA patterns generated by *35S:GFP-GUS* and *35S:GFP-EVDint/ter-GUS*. High and low molecular-weight RNA analysis using a GFP or GUS probe in two independent transgenic lines from each construct in the WT or rdr6 background. mRNA isoforms are indicated with arrows and correspond to the transcripts depicted in Fig.2A. EtBr staining of the agarose gel and miR171 probe serve as loading control for mRNAs and sRNAs, respectively. **(B)** Western analysis of the translation products from *GFP* and *GFP-GUS* transcripts. Coomassie (coom.) staining as a loading control. **(C)** Nuleo-cytosolic distribution of *35S:EVD* and *35S:GFP-EVDint/ter-GUS* RNA isoforms in *rdr6* relative to that of *ACT2* analyzed by qPCR. RNA extracted from Total, nuclear (Nucl) and cytoplasmic (Cyto) fractions was reverse transcribed with random hexamers and oligo(dT). snoRNA U5 is shown as a nuclear-only RNA control. **(D)** Same as in (C) but using exclusively oligo(dT) to reverse transcribe poly(A)+ RNAs. Both in (C) and (D), qPCR was performed on n=3 biological replicates; bars: standard error. (*) = p-value < 0.05, (**) = p-value < 0.01 (two-sided t-test between indicated samples).

### Intron retention causes selective nuclear seclusion of *flGAG-POL*-like mRNAs

What mechanism linked to splicing-coupled PCPA might underpin the differential translation of *shGAG*-like versus *flGAG-POL*-like mRNAs? Noteworthy, splicing generally enhances mRNA nuclear export and translation (Valencia *et al*, 2008; Sørensen *et al*, 2017). Conversely, polyadenylated, unspliced mRNAs are retained in the nucleus in Arabidopsis and only exported to the cytoplasm upon splicing (Jia *et al*, 2020). Moreover, 5’ splice motifs and U1 snRNP binding promote chromatin tethering of long non-coding RNAs in animal cells (Lee *et al*, 2015; Yin *et al*, 2020). We thus tested if intron-retention might promote nuclear sequestration of the unspliced *flGFP-GUS* and *flGAG-POL* or if, conversely, splicing might favor export of *shGFP* and *shGAG* to the cytoplasm, thereby selectively promoting their translation. We performed nucleo-cytosolic fractionation (Fig.S5C) to analyze the relative distributions of *EVD*-derived RNA isoforms produced in *3S:EVD_wt_* or *35S:GFP*-*EVD_int/ter_*-*GUS* plants, using spliced/unspliced isoform-specific PCR amplification. Additionally, unspliced isoforms were selectively analyzed using qPCR primer sets designed to amplify sequences located near the 3’ end of *flGAG-POL* or *flGFP-GUS,* and absent from *shGAG* and *shGFP* (Fig.1A, Fig.2A). A similar approach was used to differentiate the unspliced versus spliced *ACTIN* mRNA (Fig.S5D). Finally, the nuclear-only snoRNA U5 (Fig.5C) was used as a control to assess the quality of nuclear enrichments. To optimize accumulation of both types of RNA isoforms, the experiments were all conducted in the PTGS-deficient *rdr6* background.

The analysis revealed strikingly distinct nucleo-cytosolic distribution patterns for the full-length *versus* short spliced mRNAs from both systems. Indeed, while the spliced *shGFP* and *shGAG* were found predominantly in the cytosol (Fig.5C), *flGAG-POL* and *flGFP-GUS* were strongly enriched in nuclear fractions (Fig.5C, Fig.S5D). To validate that nuclear unspliced full-length transcripts are *bona fide* poly(A)^+^ mRNAs as opposed to nascent transcripts or splicing intermediates, cDNA from the same RNA samples was synthesized using exclusively oligo-dT to capture polyadenylated RNAs only. This approach generated comparable results (Fig.5D), indicating that nuclear full-length transcripts are properly terminated mRNAs. Corresponding results were obtained in *epi15* F11 plants displaying endogenous *EVD* reactivation (Fig.S5E-F). Collectively, these findings suggest that the unique splicing behavior of *EVD –* which is recapitulated in *GFP*-*EVD_int/ter_*-*GUS –* not only allows production of the GAG-encoding *shGAG* subgenomic mRNA, but simultaneously promotes nuclear retention of *flGAG-POL*. This is likely contributing to the disproportionate translation of *shGAG* over *flGAG-POL*, although we do not exclude the involvement of other processes. Under these premises, splicing-coupled PCPA likely predisposes *shGAG*, as opposed to *flGAG-POL*, to one or several co-translational processes which, in turn, signal(s) RDR6 recruitment.

### Saturation of co-translational mRNA decay unlikely triggers *shGAG* siRNA production

In plants and fungi, decapping coupled to 5’->3’ exonucleolytic activity operated by cytosolic XRN proteins regulate the intrinsic half-life of most actively translated transcripts by degrading decapped mRNAs after the last translating ribosome (Kastenmayer & Green, 2000; Hu *et al*, 2009; Pelechano *et al*, 2015; Yu *et al*, 2016). Of the three Arabidopsis XRNs, XRN2 and XRN3 are nuclear, whereas XRN4 is cytosolic and, hence, mediates co-translational mRNA decay (Gregory *et al*, 2008; Kurihara, 2017; Yu *et al*, 2016). Remarkably, transcripts undergoing improper decapping and/or XRN4-mediated exonucleolysis constitute competing substrates for RDR6 in Arabidopsis (Gazzani, 2004; Gy *et al*, 2007a; Gregory *et al*, 2008; Moreno *et al*, 2013; Martínez de Alba *et al*, 2015) (Fig.6A). For instance, loss-of-RDR6 function suppresses the lethality of decapping mutants by preventing production of undesirable siRNAs from hundreds of endogenous mRNAs (Martínez de Alba *et al*, 2015). Conversely, loss of XRN4 activity enhances RDR6-dependent PTGS (Gy *et al*, 2007a; Gregory *et al*, 2008; Moreno *et al*, 2013). These observations strongly suggest that RDR6-dependent PTGS takes over co-translational mRNA decay when this process becomes saturated by highly abundant and/or highly translated mRNAs.

**Figure 6.**
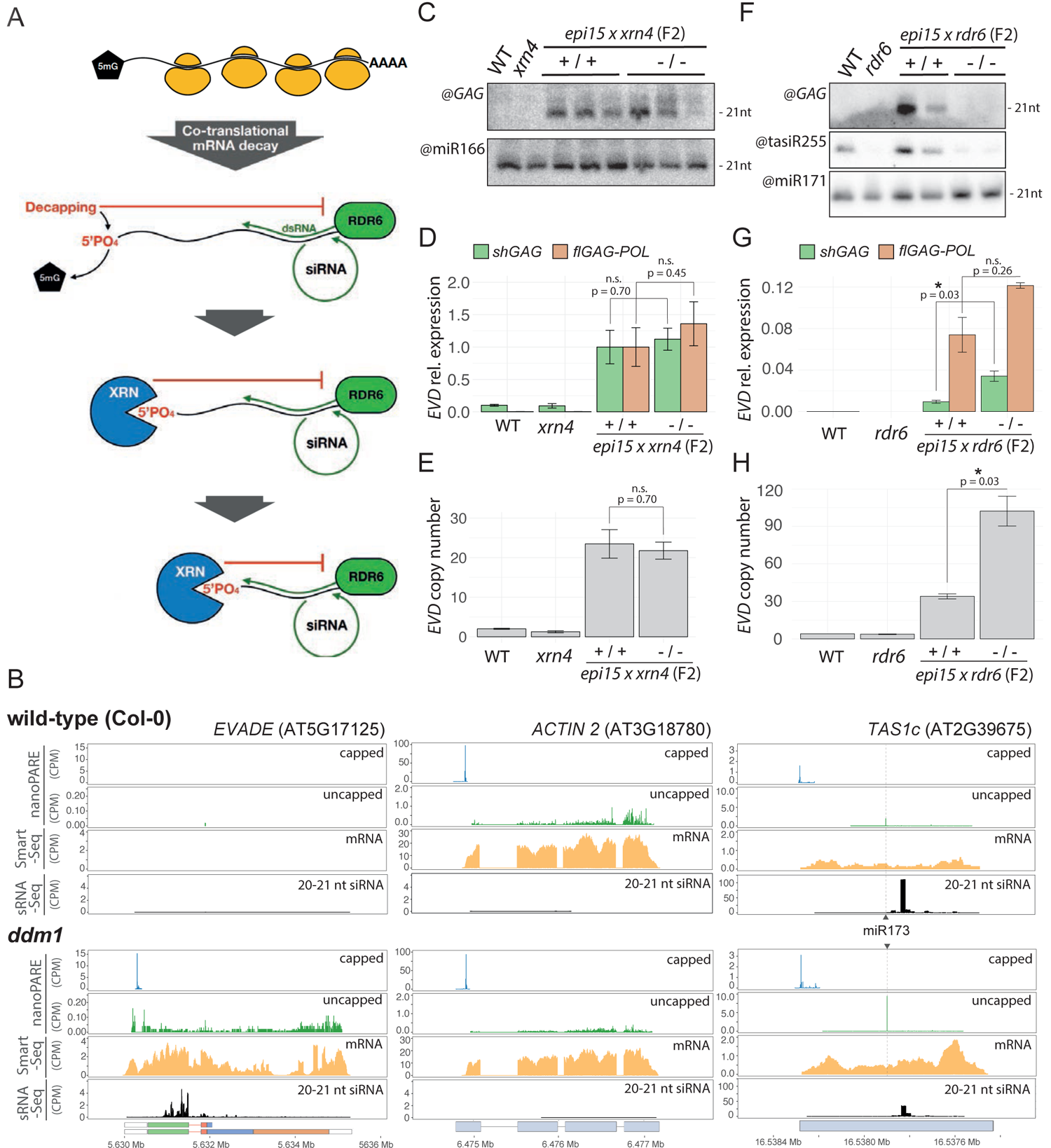
XRN4-mediated co-translational mRNA decay does not influence siRNA production from shGAG. **(A)** During co-translational decay, mRNA turnover is initiated by decapping of actively translated mRNAs. This exposes 5’ monophosphate (5’PO4) groups required for the 5’->3’ exonucleolytic activity of XRNs. As evidenced by the RDR6-dependent production of siRNAs from dozens of endogenous loci in decapping and *xrn4* mutants of Arabidopsis, plant mRNAs engaged in co-translational decay can become substrates for siRNA biogenesis if they are not degraded by, or their levels saturate, this process. While depicted here, for simplicity, as a co-translational event, RDR6 action onto such RNA likely occurs in specialized“siRNA bodies” adjacent to P-bodies **(B)** *EVD*, *ACT2* and *TAS1c* capped and uncapped 5’ends from nanoPARE and Smart-seq2 libraries along 20-21 nt siRNA in *WT* and *ddm1*. **(C-E)** *EVD* genomic proliferation in homozygous *xrn4* mutant and WT backgrounds in F2 plants from a cross between *xrn4* and epi15 exhibiting active *EVD* mobilization. (C) sRNA blot analysis using an anti-GAG probe. (D) Relative expression levels of *shGAG* and *f/GAGPOL* normalized to *ACT2* and to AT4G26410 levels. (E) EVD genomic copy number quantification by qPCR. qPCR analysis was performed on three biological replicates for controls and the three independent *WT* and mutant F2 lines displayed in B. **(F-H)** Same as (C-E) but with the rdr6 versus *WT* epi15 backgrounds. Analysis was performed in the two independent *WT* and mutant F2 lines displayed in E. In all qPCR panels: error bars display standard errors. (ns.) = non-significant, (*) = p-value < 0.05, (**) = p-value < 0.01 (two-sided t-test between indicated samples)

Likewise, we reasoned that intense translation might overwhelm XRN4-mediated co-translational decay of *shGAG* and thereby concurrently promote RDR6 action (Fig.6A). This would predict an accumulation of RNA degradation fragments (reflecting XRN4 activity) coinciding with siRNA accumulation. PARE (parallel amplification of RNA ends) and related methods map mostly XRN4 products associated with co-translational decay as well as non-translational RNA cleavage events, *e.g.* miRNA-mediated slicing of non-coding RNAs (Gregory *et al*, 2008; Schon *et al*, 2018). We therefore conducted nanoPARE analyses, which capture both capped and uncapped RNA fragments (Schon *et al*, 2018), in *ddm1 vs* WT Arabidopsis (Fig.S6A). Simultaneously, mRNA-seq (*i.e.* SMART-seq2) was conducted on the same RNA to monitor gene expression (Schon *et al*, 2018). Analysis of *TAS1c*, which undergoes miR173-mediated slicing, confirmed that the ensuing 3’ RNA cleavage fragment, a common substrate of XRN4 (Schon *et al*, 2018), was readily detected in both backgrounds, despite spawning vast amounts of RDR6-dependent siRNAs (Fig.6B). Analyzing *EVD* upon its reactivation in *ddm1* revealed a low level of RNA degradation fragments spanning the entirety of *EVD* despite the siRNAs being exclusively derived from *shGAG*. Had RNA degradation contributed to siRNA biogenesis, these species would be expected to be distributed along the entirety of *EVD,* encompassing both *shGAG* and *flGAG-POL*. Inspection of the housekeeping *ACT2* locus revealed a similar ORF-spanning degradation pattern, albeit at substantially higher levels (∼10-folds), presumably reflecting the higher transcript abundance. However, *ACT2* does not spawn siRNAs (Fig. 6B). These observations therefore reveal no overt correlation between abundance of RNA degradation products, siRNA production and/or polysome association.

The above results did not formally exclude the possibility that at least some *EVD-* associated degradation products identified by nanoPARE might contribute to siRNA biogenesis *via* competing RDR6 *vs* XRN4 activities. This would be genetically diagnosed by an increased accumulation of *shGAG* siRNAs in *xrn4* in contrast to their loss in *rdr6* (Gy *et al*, 2007a; Gregory *et al*, 2008). To test this idea without the potential complication of *EVD* overexpression artificially saturating XRN4 activity in *35S:EVD_wt_*, we introgressed the *xrn4* null-mutation into *epi15* at the early F8 inbred generation, when PTGS of *EVD* is commonly initiated (Marí Ordóñez *et al*, 2013). As negative controls, we used loss-of-function alleles of nuclear XRN2 and XRN3, which, by not contributing to co-translational mRNA decay, should not influence siRNA production. Finally, the *rdr6* mutation was introgressed in parallel, to prevent *shGAG* siRNA biogenesis. We analyzed two-to-three independent lineages with WT *versus* homozygous mutant backgrounds isolated from segregating F2s. However, neither *xrn4* nor *xrn2/xrn3* differed from the WT background with regard to *EVD* expression, copy number, or *shGAG* siRNA levels (Fig.6C-E, Fig.S6B-G). In contrast, *EVD* expression and copy numbers were increased in *rdr6*, coinciding with reduced *shGAG* siRNA levels (Fig.6F-H). We conclude from these collective results that saturation of XRN4-dependent co-translational mRNA decay (Fig.6A) is unlikely to underlie *shGAG* siRNA production. siRNAs are, instead, abruptly spawned from the middle up to the 3’ end of *shGAG*, as if their production coincided with a discrete co-translational event (Fig.6A). A similar rationale should apply to the discrete *shGFP*-centric siRNA pattern spawned from *GFP*-*EVD_int/ter_*-*GUS* (Fig.2C).

### The initiation of RDR6 activity coincides with isolated and intense ribosome stalling events

To overcome the caveat of *EVD* cell-specific expression (Marí Ordóñez *et al*, 2013) and simultaneously investigate which co-translational event(s) might trigger the siRNA patterns in both *shGAG* and *shGFP*, we generated RIBO-seq datasets (Ingolia *et al*, 2009) from *35S:EVD_wt_* and *35S:GFP*-*EVD_int/ter_*-*GUS*. This resulted in high-quality ribosome footprints (RFPs) displaying the characteristic triplet periodicity (Fig.S7). We found that *EVD* RFPs in the *35S:EVD_wt_* background map near-exclusively onto *shGAG*, underscoring its preferential translation (Fig.4A, 7A). However, a strong and isolated footprint peak was detected near the middle of the *shGAG* ORF (Fig.7A), suggesting intense ribosome stalling at this position. This stalling peak was also found within the *shGAG* coding sequence of endogenous *EVD* in one public *ddm1* RIBO-seq library (Kim *et al*, 2021) (Fig.S8A). By specifying codon occupancy of ribosome P-sites – the sites of peptidyl transfer activity – reflecting the codon dwell time, we found that >35% of *shGAG* translating ribosomes are located on two consecutive codons (pos.148-149) coinciding with this peak (Fig.7B). Having normalized these proportions to ORF lengths, we compared them to those of actively translated Arabidopsis mRNAs. To exclude artefacts from transcripts with low coverage, we restricted our analysis to the most abundant mRNA isoforms with coverage available for more than 70% of ORFs, as described (Sabi & Tuller, 2015). We found that *shGAG* ranks among the top 4.01 and 2.77 % (in WT and *rdr6* backgrounds respectively) of Arabidopsis transcripts displaying the most intense stalling events (Fig.7C). Remarkably, overlaying siRNAs and codon coverage intensity revealed that the intense stalling position coincides nearly exactly with the 5’ starting point of the RDR6-dependent *EVD* siRNA pattern (Fig.7D). Stalling is likely causal, not a consequence of RDR6 recruitment, because it also occurs in the *rdr6* background (Fig.7A).

To explore further a possible link between discrete, intense ribosome stalling and RDR6 recruitment, RFPs were conducted in the *35S:GFP*-*EVD_int/ter_*-*GUS* background. As seen above for *shGAG* versus *flGAG-POL* in the *EVD* context, the analysis confirmed the vastly disproportional translation of *shGFP* versus *flGFP-GUS* (Fig.S8B). It also identified a major stalling site only in the *shGFP* ORF (whose detection was enhanced in the *rdr6* background) in which two prominently covered and consecutive codons (pos. 235-236) accounted for ∼40% of footprints (Fig.S8C). Similarly to *shGAG*, *shGFP* ranked among the top 3.21 to 4.14% Arabidopsis transcripts displaying the most intense stalling events (Fig.S8D). Furthermore, this stalling site was located between major peaks of *shGFP* siRNAs, in this case, in both the 5’ and 3’ directions within *GFP*-*EVD_int/ter_*-*GUS* (Fig.S8E).

A recent model advocates a possible link between suboptimal codon usage and PTGS initiation in plants (Kim *et al*, 2021). However, overlaying the Arabidopsis codon adaptation index with the codon coverage of ribosomes on *shGAG* and *shGFP* did not reveal any overt correlation between ribosome stalling and codon suboptimality (Fig.S9). The cited study showed that codon optimization in a region corresponding, surprisingly, to the *shGAG* 3’UTR enhanced translation of a linked luciferase ORF (Kim *et al*, 2021). Yet, our analysis shows that neither CG nor CG3 content (CG content on the 3^rd^ codon position) overtly influences ribosome association along *shGAG* or *shGFP*, let alone the intense stalling event detected on either mRNA (Fig.S10). Two consecutive codons at the stalling site identified on *shGAG* code for proline and glycine (Fig.S9A) and, interestingly, single prolines and/or glycines at P-sites correlate with ribosome stalling in animals and fungi (Artieri & Fraser, 2014; Sabi & Tuller, 2015; Zhao *et al*, 2021). By contrast, the >45% codon occupancy on *shGFP* occurs on unrelated, consecutive glutamate and leucine codons (Fig.S9B). In addition to the identity of some codons, secondary RNA structures have been correlated with ribosome stalling (Doma & Parker, 2006; Yan *et al*, 2015; Bao *et al*, 2020), including G-quadruplexes (Song *et al*, 2016; Fay *et al*, 2017). In particular, sites of “ribothrypsis” – a ribosome stalling-induced process recently described in metazoans – are positively correlated with such occurrences within ORFs (Ibrahim *et al*, 2018). By forming secondary structures, G-quadruplexes are thought to act as “roadblocks” hampering proper ribosome progression during elongation (Song *et al*, 2016). While G-quadruplex scoring along the *shGAG* and *shGFP* mRNA did reveal potential hot spots of such motifs, they were localized far upstream or downstream of each identified ribosome stalling site (Fig.S9). Overall, *EVD* and *GFP*-*EVD_int/ter_*-*GUS* transcripts display similar behavior, whereby RDR6-dependent production of *shGAG*- or *shGFP-*only siRNAs coincides with highly localized and unusually intense ribosome stalling events. While stalling events have likely distinct causes for each transcript, they nonetheless appear to stimulate co-translational processing of RNA intermediates that, in turn, serve as RDR6 substrates.

### Ribosome stalling correlates with production of 5’-hydroxy 3’-cleavage fragments that possibly serve as RDR6 substrates

As described above, nanoPARE in *ddm1* did not reveal any discrete RNA products with 5’ ends mapping consistently at, or near, the stalling site in *shGAG*. We also failed to detect such products using classic 5’ RACE (Llave *et al*, 2002). Noteworthy, this technique relies on a 5’ monophosphate (5’P) for RNA ligation of 5’ adaptors (Silber *et al*, 1972; Wang & Fang, 2015). Intriguingly, 5’P was reported to be absent from various 3’ cleavage RNA fragments produced co-translationally in budding yeast, including upon ribosome stalling (Peach *et al*, 2015; Navickas *et al*, 2020). A lack of 5’P is also strongly suspected for the 3’ cleavage products of ribothrypsis (Ibrahim *et al*, 2018). Since siRNA production from *EVD* initiates just downstream of the major stalling site (codons 148-149), we thus considered the possibility that discrete 3’ cleavage RNA fragments devoid of a 5’P – and thus akin to the 5‘OH RNA associated with the above-mentioned processes (Peach *et al*, 2015; Navickas *et al*, 2020; Ibrahim *et al*, 2018; D’Orazio *et al*, 2019) – might constitute RDR6 templates (Fig.7E).

To explore such a connection and simultaneously characterize and map the 5’ ends of putative *shGAG* 3’ cleavage fragments, we used the RtcB RNA ligase. RtcB contributes to tRNAs splicing by ligating RNAs with 3’P ends (or 2’,3’-cyclic phosphate) to 5’OH ends, and was used previously to map co-translational RNA cleavage fragments in yeast (Desai & Raines, 2012; Peach *et al*, 2015). A 5’ RNA adaptor with a 3’P end was therefore RtcB-ligated to total RNA extracted from plants expressing *35S:EVD* or non-transgenic controls, both in the *rdr6* background. Use of *rdr6* prevented conversion of potential RDR6 templates into dsRNA as well as the accumulation of confounding cleavage fragments potentially caused by the ensuing secondary siRNAs. The ligated RNA was then subjected to reverse transcription using *EVD*-specific primers surrounding the major stalling site (Fig. 7F; Region #1), amplified through PCR, and cloned following standard RACE procedures. Based on the *EVD* ribosome footprint profile (Fig. 7B), we also investigated two additional regions more covered with ribosomes than expected (Fig. 7F; Regions #2 & 3). Only region #1 yielded detectable amplification products within the expected size range. Nonetheless, gel excision within the anticipated size ranges followed by cloning was performed for all regions in all genotypes (Fig. S10A-C). Sanger sequencing revealed that 30 out of 36 fragments cloned from region #1 displayed 5’OH ends consistently mapping at nucleotides 447-448, strikingly defining the intense ribosome stalling site on *shGAG* (Fig. 7F, S10D) from which siRNA production is initiated (Fig. 7D). By contrast, the clones obtained from regions #2 and #3 were either devoid of *EVD* sequences or empty. These results are consistent with the notion that the intense ribosome stalling event correlates with breakage of the *shGAG* RNA, and that the ensuing 5’OH fragments serve as templates for RDR6 to initiate dsRNA production and downstream siRNA processing. Given that XRNs require a 5-P for their 5’->3’ exonucleolytic activities (Stevens, 2001; Schon *et al*, 2018), this could explain the insensitivity of *shGAG*-derived siRNA accumulation to any *xrn* mutation and to *xrn4* in particular (Fig.6). Being linked to 3’ cleavage fragments inaccessible to the competing activity of XRN4, ribosome stalling might thus optimize the recruitment of RDR6 on *shGAG* for PTGS initiation. We note that nanoPARE, while being indiscriminative of RNA 5’-ends (including 5’OH) requires a 3’ polyA tail to generate cDNA. Thus, the fact that the technique failed to detect the *shGAG* 5’OH cleavage fragments could indicate that they are indeed mostly poly(A)^-^. Preliminary PAGE-based analyses as conducted for the 3’ ends of ribothrypsis products in mammalian cells (Ibrahim et al. 2018) suggested that discrete *shGAG*-derived 3’ cleavage fragments are found in the poly(A)^-^ fraction isolated from *35S:EVD* in the *rdr6* background (Fig.S11A). Lack-of-poly(A) could further optimize RDR6 recruitment because the enzyme is inhibited *in vitro* by 3’ adenosine stretches (Baeg *et al*, 2017).

**Figure 7.**
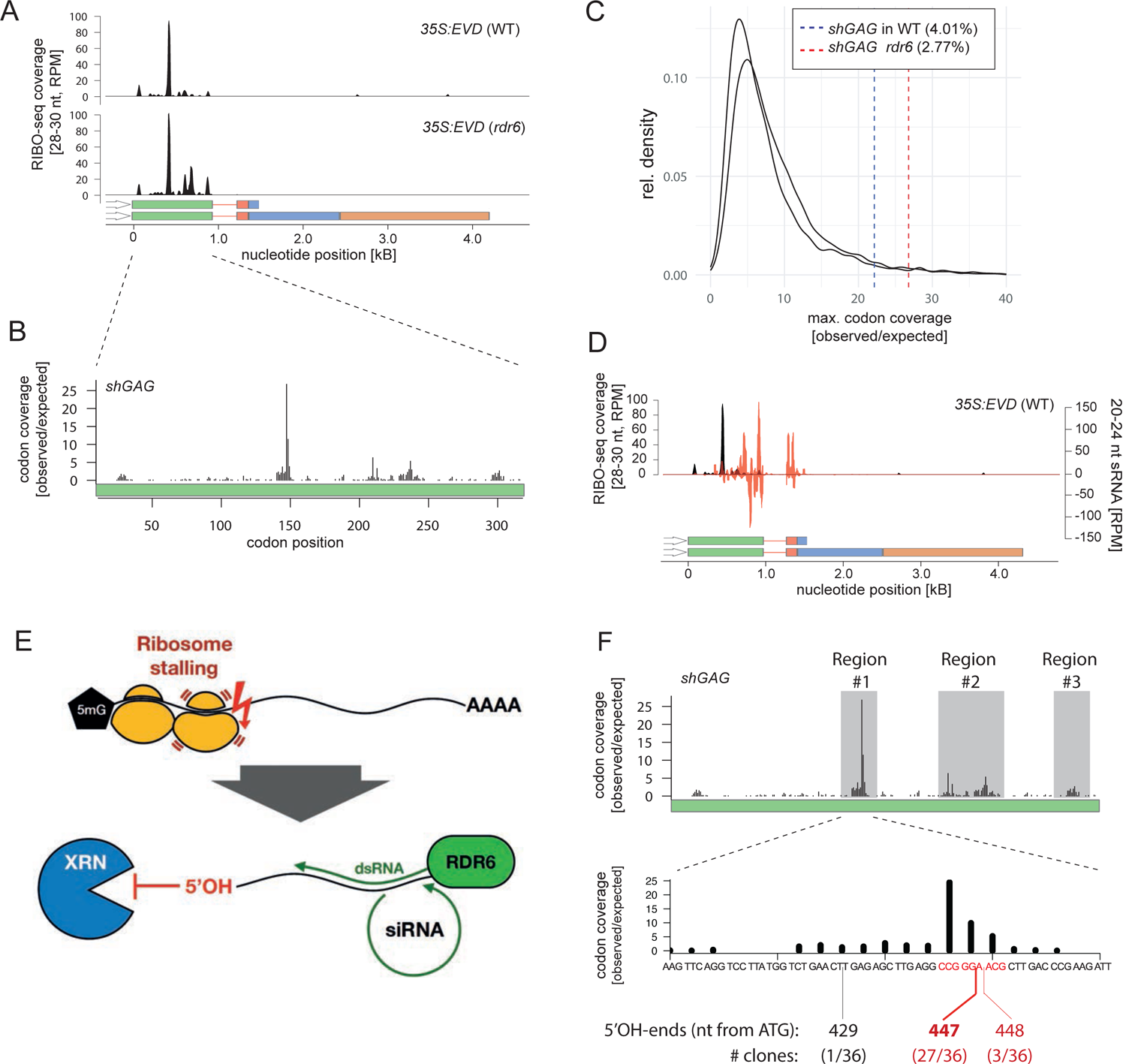
Intense, discrete ribosome stalling on shGAG correlates with RDR6-dependent siRNA accumulation. **(A)** RIBO-seq coverage profiles from *35S:EVD* in *WT* or *rdr6*. RPM: Reads per million. *(B)* Ribosomal footprints on *shGAG* in *rdr6* displaying codon occupancy at P-sites to calculate codon coverage. The coverage observed at each codon position was divided by the expected mean coverage along the entire GAG coding sequence. *(C)* Maximal individual codon coverage over the expected coverage for all translated transcripts of Arabidopsis. Vertical lines indicate the strength of stalling sites of *shGAG* in the *WT* or *rdr6* background. Percentages specify the proportion of transcripts with more pronounced stalling events than *shGAG*. **(D)** Overlay between *35S:EVD* siRNAs (red) and RIBO-seq profiles (black) in the *WT* background. **(E)** Schematic representation of putative ribosome stalling-linked mRNA breakage generating 5’OH ends. Lack of 5’PO4 prevents XRN 5’->3’ exonucleolytic activity (see Fig.6A), granting the RNA to be used as template by RDR6. **(F)** Overlap between ribosomal footprints and mapping of 5’OH ends from *35S:EVD* in *rdr6* cloned through RtbC ligation. Regions investigated are highlighted in grey. 5’OH ends were only successfully cloned from region #1. Alignment of sequenced clones to *EVD* is displayed in Fig.S9.

## DISCUSSION

### Translation as an initiator of PTGS and epigenetic silencing

Protein synthesis is commonly merely seen as a target of PTGS by reducing the amount of available RNA and/or interfering with translation. Our study identifies translation also as a trigger of PTGS. This became evident after epigenetic reactivation of *EVD*, from which splicing-coupled PCPA generates separate RNA isoforms from a single transcription unit. Of the two, the shorter subgenomic *shGAG* RNA undergoes disproportionate translation over *flGAG-POL* as an indispensable feature of *Ty1/Copia* biology because this likely provides the stochiometric protein balance necessary for efficient amplification and mobilization of the element. This process, however, concomitantly stimulates RDR6 activity. *shGAG* translation efficacy *per se* is within the range of moderately translated Arabidopsis mRNAs and is unlikely to explain this effect, nor do *GAG* expression or abundance. Rather, an exceptionally intense and highly discrete ribosome stalling event predisposes *shGAG* to RDR6-dependent PTGS. Our data also suggest how intron-retention in combination with active splicing accounts for the mostly nuclear *versus* cytosolic localization of *flGAG-POL versus shGAG*, respectively. Their asymmetrical subcellular distribution concurrently rationalizes (*i*) the disproportionate translation efficacies of each mRNA, (*ii*) the *shGAG*-centric distribution of translation-dependent *EVD*-derived siRNAs and, consequently, (*iii*) the contrasted sensitivity of each isoform to cytosolic PTGS. Splicing-coupled PCPA probably underlies most, if not all, of features *i-iii* because they were recapitulated with the *GFP*-*EVD_int/ter_*-*GUS* construct containing the *shGAG* intron and proximal PCPA signal (Fig.2, S4 and 5). Since splicing-coupled PCPA is at the very core of the *Ty1/Copia* genome expression strategy (Oberlin *et al*, 2017), the process described here for *EVD* is likely to be broadly applicable.

Being mostly nuclear, *flGAG-POL*, the template for RT required for mobilization, is neither a potent trigger nor a target of PTGS, likely explaining why increasing amounts of *shGAG* siRNAs have little impact on *EVD’s* genomic proliferation over successive epi15 inbred generations (Marí Ordóñez *et al*, 2013). Previously attributed to GAG-mediated protection of *flGAG-POL* as part of VLPs (Marí Ordóñez *et al*, 2013), we now consider *flGAG-POL* nuclear retention as an additional and perhaps major contributor to this shielding effect. The ensuing rise in *EVD* genomic copies causes increasing levels of RDR6-dependent *shGAG* dsRNA over generations. We previously suggested that these levels eventually saturate DCL4/DCL2 activities in the highly cell-specific expression domain of *EVD*, acting as a prerequisite to DCL3 recruitment and RdDM, ultimately causing LTR methylation and TGS of all *EVD* copies (Marí Ordóñez *et al*, 2013). This proposed saturation-coupled PTGS-to-TGS switch invariably occurs in epi15 and other *EVD*-reactivating epiRILs when the *EVD* copy number reaches 40-50 (Marí Ordóñez *et al*, 2013), causing only sporadic and minor developmental defects even in advanced generations (Marí Ordóñez *et al*, 2013; Mirouze *et al*, 2009; Quadrana *et al*, 2016). By contrast, *EVD* copy number increases well beyond 80 in *rdr6* mutants already in F2s (Fig.6H) displaying loss-of-fertility (Fig.S11B) likely solely ascribable to enhanced *EVD* proliferation. These data attests to a central role for RDR6 in controlling *EVD*’s mobilization and perhaps that of other autonomous TEs, at the level of translation. At least in the multi-generational context of epi15, our results also establish a *hitherto* unrecognized role for translation as an initiator of not only PTGS, but also, ultimately, of epigenetic silencing and TGS.

### Translation-dependent silencing as a sensor for *de novo* invading, foreign genetic elements

The vast majority of *ddm1*-reactivated TEs that spawn RDR6-dependent siRNAs is composed of *LTR/Gypsy* elements (Fig.4), which is the family most prominently associated with easiRNA production (Creasey *et al*, 2014; Borges *et al*, 2018). Arabidopsis *LTR/Gypsy* elements generally display significantly shorter-than-full-length ORFs as compared to the other main classes of Arabidopsis TEs, including the *LTR/Copia* family to which *EVD* belongs (Oberlin *et al*, 2017) (Fig.4C). Although they likely constitute, therefore, degenerated transcription units, a substantial fraction of such *LTR/Gypsy* is nonetheless highly expressed as a possible source of abundant aberrant RNAs (Fig.4B). Thus, alternatively or concurrently to easiRNA production, some of these *LTR/Gypsy* remnants might also enter the RDR6 pathway by saturating RQC either co-transcriptionally or post-transcriptionally. While this process possibly underlies a previously documented expression-dependent form of innate TE silencing (Panda *et al*, 2016; Fultz & Slotkin, 2017), such loci might in turn autonomously produce siRNAs and become sources of identity-based silencing. Regardless, the combined action of all these silencing pathways likely explains why most siRNA-generating TEs in *ddm1* do not actively engage translation, as evidenced by their conspicuous underrepresentation on polysomes (Fig.4A; quartile 4) (Oberlin *et al*, 2017). In fact, siRNA-generating TEs are equally or even less polysome-associated than are non-coding RNAs (Fig.4A), indicating that there is no general correlation between siRNA production and translation. Conversely, numerous TEs are translated, yet do not spawn siRNAs (Fig.4A). These data contradict recent claims advocating a general correlation between siRNA production and translation based on the untested premise that most siRNA-generating TEs are translated (Kim *et al*, 2021), whereas they are, in fact, absent from polysomes (Oberlin *et al*, 2017) (Fig.4A). Based on our experimental findings we argue, on the contrary, that the process of “translation-dependent silencing” (TdS) described here is an attribute of only a handful of evolutionary young TEs. These chiefly include *EVD,* which concurrently undergoes productive translation (mostly of *shGAG*) and spawns RDR6-dependent siRNAs (Figs.1, 4 and S1).

*EVD* is among the few autonomously transposing LTR/TEs in the Arabidopsis *Col-0* genome (Mirouze *et al*, 2009; Tsukahara *et al*, 2010; Gilly *et al*, 2014; Reinders *et al*, 2009a) and, as such, is unlikely controlled by identity-based mechanisms. TdS might enable the plant to detect its activity as the first line of defense against *de novo* invasions, for instance upon horizontal transfer of active TEs. TdS may likewise underpin silencing triggered upon experimental transfer of “exogenous” TEs between species separated by millions of years of evolution (Hirochika *et al*, 2000; Fultz & Slotkin, 2017). Viruses divert a substantial fraction of the host translational apparatus to their highly compact and TE-like genomic and subgenomic RNAs (GAO, 2003; Dreher & Miller, 2006; Sztuba-Solińska *et al*, 2011), which might also predispose them to TdS. In all these circumstances, a key feature of TdS is an innate ability to detect transcripts by virtue of their foreign – as opposed to aberrant – nature, independently of any sequence homology to the host genome. We propose that foreignness is perceived by anomalies manifested during active translation, which likely include abnormal ribosome stalling, as discussed below.

### Possible cause(s) of ribosome stalling

Our findings raise the key question of what molecular signal(s) might cause the unusually intense stalling events observed in the s*hGAG* and *shGFP* ORFs that possibly activate TdS. While studies of ribosome stalling in plants are very scarce, suboptimal codon usage has been widely considered as one of its possible causes (Rocha, 2004; Quax *et al*, 2015). Indeed, extensive stretches of rare codons trigger ribosome stalling and RNA degradation when artificially engineered in reporter mRNAs (Li *et al*, 2006; Yang *et al*, 2019; Park & Subramaniam, 2019). However, it has been noted that this rarely applies to endogenous mRNAs, where non-optimal codons are common and play roles in translation regulation (Hanson & Coller, 2018; Carneiro *et al*, 2019). Accordingly, recent in-depth analyses in animals and fungi indicate that codon usage *per se* is not predictive of stalling on endogenous mRNAs, and that other factors might modulate ribosome dwell times (Gardin *et al*, 2014; Dana & Tuller, 2014; Rodnina, 2016; Zhang *et al*, 2017; Wu *et al*, 2019). *shGAG* and *shGFP* are devoid of rare codons among other inspected features, suggesting that no universal sequence/structure feature might underpin the stalling events correlating with TdS. Rather, mRNA-intrinsic and variable features, or combinations thereof, might be involved. mRNA-extrinsic features could also contribute to intense and discrete stalling in *shGAG* via *trans*-interactions involving specific RNA sequence-motifs and RNA-binding proteins (RBP), a circumstance that can hinder ribosome progression (Babitzke *et al*, 2009; Iwakawa & Tomari, 2015; Zhang *et al*, 2017). The action of AGO-miRNA complexes could illustrate how RBPs may, under some circumstances, elicit TdS, for instance during the intricate biogenesis of *TAS3* trans-acting (ta)siRNAs (Hou *et al*, 2016; Xia *et al*, 2017). We note that a recent model proposed by Kim and coworkers (Kim *et al*, 2021) contends that PTGS via RDR6 might be caused by a multitude of presumptive translation stalling events. These were allegedly ascribed to pervasive suboptimal codons and low content in CG and CG3 along the ORFs of certain mRNAs, including TE-derived RNAs. However, this interpretation is not compatible with the highly discrete nature of the stalling events experimentally detected in our study and the noticeable absence from polysomes of the TE RNAs used to build this model, *EVD* excepted (Fig.4A).

### Possible mechanism(s) of TdS

Intense stalling is usually resolved on the protein side by ubiquitin-mediated proteolysis (Joazeiro, 2019). On the RNA side, it is released by translation-decoupled RNA degradation related to, albeit distinct from, co-translational decay (Ikeuchi *et al*, 2018). One such process is XRN-mediated exonucleolysis operating in dedicated processing (P) bodies (Maldonado-Bonilla, 2014), yet 5’OH mRNA fragments are not directly accessible to XRN action, which requires 5-P *termini* (Stevens, 2001). In budding yeast, the Trl1 kinase progressively phosphorylates these fragments to license their degradation by XRNs (Navickas *et al*, 2020), a process also likely occurring during ribothrypsis in mammalian cells (D’Orazio *et al*, 2019). Alternatively, or additively, the mouse and fission yeast DXO/Rai1, which removes incomplete 5’-mRNA caps, catalyzes the removal of 5’OH ends, exposing 5’P for subsequent 5’->3’ exoribonuclease activity (Doamekpor *et al*, 2020). An Arabidopsis DXO1 catalytic mutant for cap surveillance also displays such an activity in vitro, although it shows higher affinity for 5’P substrates (Doamekpor *et al*, 2020). Whether AtDXO1 antagonizes TdS in vivo will have to be further investigated.

In contrast to RNAi-deficient budding yeast or RNAi-proficient mammalian cells, plants display RDR activities (Stein *et al*, 2003; Drinnenberg *et al*, 2009). We suggest that in these organisms, 5’OH *termini* would not only disqualify XRN4 action, but concurrently optimize that of RDR6, which is known to compete with XRN4 for substrates, including those evading co-translational decay (Gy *et al*, 2007b; Gregory *et al*, 2008). Incidentally, 5’P-*termini* are also generated during mRNA decapping (Hu *et al*, 2009), potentially explaining, for similar reasons, why decapping Arabidopsis mutants are hypersensitive to RDR6 (Martínez de Alba *et al*, 2015). RDR6 action in TdS is possibly further facilitated by the striking physical proximity of P-bodies – where unresolved 5’OH RNA fragments should primarily accumulate – with the so-called “siRNA bodies” (Martínez de Alba *et al*, 2015). RDR6 and co-factors congregate in these bodies for tasiRNA processing, which, we suggest, are also the sites of *shGAG* siRNA biogenesis. We emphasize that converting the 5’OH-RNA fragments into dsRNA would lead to siRNA production accompanied by limited exonucleolytic degradability of RNA. Widespread RNA degradation intermediates spanning entire transcriptional units were recently proposed to represent RDR6 substrates for translation-coupled PTGS including in the case of *EVD* (Kim *et al*, 2021). This model, however, does not explain the discrete siRNA pattern observed exclusively on *shGAG* yet not *flGAG-POL*. It neither explains why highly abundant, ORF-spanning RNA degradation products, such as those derived from *ACT2,* do not correlate with siRNA production.

Capped but poly(A)^-^ 5’ RNA fragments are expected to be concurrently produced during the RNA breakage process, that should also be amenable to RDR6 activity. Yet, only a small siRNA peak is detected just upstream of the intense stalling site on *shGAG*. Unlike the XRN4-protected 5’OH 3’ fragments, however, these 5’ fragments would be *a priori* devoid of features preventing their SKI2-mediated exosomal 3’-5’ degradation, which indeed can outcompete RDR6 action (Zhang *et al*, 2015; Branscheid *et al*, 2015). Differential competition between SKI2-mediated degradation *versus* RDR6 activity, as previously reported for some miRNA targets (Branscheid *et al*, 2015), might explain why the siRNAs pattern from *shGFP* spans both the 5’ and 3’ sequences surrounding the identified stalling site whereas these are only located 3’ to the *shGAG* stalling site of (Fig.7D, S8E). In principle, RDR6 could also pick up a multitude of RNA cleavage fragments predictably produced *via* siRNA-guided cleavage of *shGAG* by AGO1/AGO2-RISCs. However, RISC-mediated slicing produces 5’P *termini* (Martinez & Tuschl, 2004), which would qualify these RNAs as XRN4-, as opposed to RDR6-, substrates, therefore unlikely contributing prominently to *shGAG* siRNA production. A final, outstanding aspect of TdS pertains to the mechanism whereby 5’OH fragments are generated. In budding yeast, the metal-independent endonuclease Cue2 was recently shown to cleave, within the colliding ribosome’s A site, mRNAs undergoing stalling-induced no-go decay, which generates 5’OH 3’ RNA fragments (D’Orazio *et al*, 2019). The mammalian homolog, N4BP2, which additionally contains a polynucleotide kinase domain, might directly couple endonucleolysis with the 5’P-dependent XRN-licensing step evoked above (D’Orazio *et al*, 2019). We failed, however, to identify a plant Cue2/ N4BP2 ortholog. We suggest that other mechanisms might underlie what we therefore conservatively refer to as “translation-linked mRNA breakage” here. *EVD* or reporters derived thereof (e.g. *GFP*-*EVD_int/ter_*-*GUS*) might provide a paradigm to elucidate, via biochemistry and forward genetics, the mechanisms of TdS in plants.

Among other processes, transient ribosome stalling is a normal and favorable feature of translation enabling proper folding of nascent peptides (Rodnina, 2016). Accordingly, many mechanisms exist to resolve such instances (Buskirk & Green, 2017) including ribothrypsis in mammalian cells, which appears to be a widespread component of ordinary translation (Ibrahim *et al*, 2018). Like ribothrypsis, the process described here might have eluded characterization for years because its products have 5’OH and possible 3’ poly(A)^-^ *termini* inaccessible to standard RNA sequencing procedures. However, while its initiation strongly resembles that of mammalian ribothrypsis, TdS is unlikely to be ubiquitous in plants, since the aforementioned RNA products, by directly engaging RDR6 for amplified siRNA production, would promote degradation of the entire mRNA pool independently of its stalled or even merely translated status. While this would be highly detrimental as a common form of endogenous gene regulation, the process seems particularly well-suited to eliminate highly proliferating foreign RNAs such as those of viruses and TEs. We suspect that TdS might also represent a yet unexplored trigger for PTGS of transgenes encoding, in particular, non-plant ORFs with suboptimal translation features.

## Author contributions

A.M.O., S.O., and O.V. conceived and designed the study. S.O. and A.M.O. performed most experiments. R.R. cloned and sequenced the *EVD* 5’OH ends, M.T. and V.B.B. investigated *EVD* copy number in *rdr6* and RNA isoform distribution in cyto-nuclear fractions. M.A.S., A.P. and M.N. performed nanoPARE. L.L. conducted seed counting. S.O. and M.A.S performed computer analyses. S.O. and M.T performed statistical analyses. A.M.O., S.O., and O.V. analyzed the data and wrote the manuscript.

## Acknowledgements

We thank members of the Voinnet and Marí-Ordóñez labs and colleagues for critical reading of the manuscript and for discussions. This work was supported by a core grant attributed to OV by the ETH-Zürich that covered the largest part of this study, including the PhD studentships of S.O. and A.M.O. under the Life Science Zürich Graduate School program. Part of the work was also supported by the NCCR RNA & Disease funded by the Swiss National Science Foundation.

## Material and methods

### Plant material and growth conditions

Plants were grown in a growth chamber on soil at 22°C for two weeks in a 12h/12h light cycle and then transferred to a 16h/8h light cycle and pools of three to five plants were sampled for inflorescence tissue. Mutant genotypes *met1-3, dcl1-11, ddm1-2* (seventh inbred generation), *hyl1-2, rdr6-12, xrn2-2, xrn3-3, xrn4-3* plants are all derived from the Col-0 ecotype (Vongs *et al*, 1993; Peragine *et al*, 2004; Vazquez *et al*, 2004; Gy *et al*, 2007b). Genotyping primers are described in Suppl. Table 1. *met1-* derived epiRIL#15 plants (*epi15*) were described previously (Reinders *et al*, 2009b; Oberlin *et al*, 2017). *35S*:*EVDwt, 35S:EVD_mU1_, 35S:EVD_Δi_, 35S:GFP-GUS* and *35S*:*GFP*-*EVD_int/ter_*-*GUS* overexpression lines were previously depicted (Marí Ordóñez *et al*, 2013; Oberlin *et al*, 2017).

### Constructs and Plasmids

All constructs are available from addgene (www.addgene.org): 35S:EVDwt (#167119), 35S:EVDmU1 (#167121), 35S:EVDΔI (#167120), 35S:GFP-GUS (#167122) and 35S:GFP-EVDint/ter-GUS (#167123) (Marí Ordóñez *et al*, 2013; Oberlin *et al*, 2017).

### Cyto-nuclear fractionations

For each sample, twice 250 mg of 3 week-old seedlings grown in ½ strength (2.2 g/L) Murashige and Skoog medium (#M0231,Duchefa Biochemie) were ground to fine powder in liquid nitrogen and homogenized in 575 μL of lysis buffer (10 mM Tris-HCl pH 7.4, 150 mM NaCl, 0.15% IGEPAL (CA-630, Merk) and 1x cOmplete protease inhibitor cocktail (Roche)). Lysates were gently mixed and incubated on ice for 10 min. before being filtered through one layer of Miracloth. 400μL from each lysate were recovered and one set aside as Total. The second set of cell lysate were gently overlaid on top of 1 mL of cold sucrose buffer (10 mM Tris-HCl pH 7.4, 150 mM NaCl, 24% sucrose and 1x cOmplete EDTA-free protease inhibitor cocktail (#04693159001, Roche)) in protein low binding 1.5 mL tubes (LoBind, Eppendorf) by slowly pipetting against the side of the tube. Samples were centrifuged at 3500x, *g* for 10 min. to separate nuclei (pellet) from cytoplasm (supernatant). Cytoplasmic fractions were cleared by centrifugation at 14000x *g* for 1 min. in a new tube and the resulting supernatant set aside. Nuclear pellets were rinsed by inverting the tube 3-5 times without disturbing the pellet with 1 mL of 1X PBS, 0.5mM EDTA. Nuclei were spin for s. at 1300x *g* before gently removing the wash solution. Nuclei pellets were resuspended by pipetting in 200 μL of nuclear lysis buffer (10 mM Tris-HCl pH 7.4, 300 mM NaCl, 7.5 mM MgCl*_2_*, 0.2 mM EDTA pH8, 1M Urea, 1% IGEPAL and 1x cOmplete protease inhibitor cocktail). For isolation of total RNA and protein from the different fractions, samples were mixed 1 volume of acid PCI (Phenol/Chloroform/Isoamyl-alcohol, #X985 Carl Roth). In addition, nuclear fractions were further homogenized after addition of PCI by passing the sample through a 21- gauge needle with a 1 mL syringe. All steps were carried on ice or centrifuged at 4°C. Buffers were freshly prepared in advance and chilled on ice before use.

### Nucleic acid and protein extractions

RNA was extracted from frozen and ground tissue with TRIzol reagent (#93289, Sigma) and precipitated with 1x vol. of cold isopropanol. For RNA extraction from cyto- nuclear fractionations, 20 μg of glycogen (#R0551, ThermoFisher) and 0.1x vol. of sodium acetate 3M pH5.2 were mixed with recovered aqueous phases after PCI before RNA precipitation with 1x vol. of cold isopropanol. DNA was extracted using the DNeasy Plant Mini Kit (#69204, Qiagen) according to manufacturer’s guidelines. Protein of frozen and ground tissue was homogenized in extraction buffer (0.7 M sucrose, 0.5 M Tris-HCl, pH 8, 5 mM EDTA, pH 8, 0.1 M NaCl, 2% β-mercaptoethanol) and cOmplete EDTA-free protease inhibitor cocktail (#04693159001, Roche). Water-saturated and Tris-buffered phenol (pH 8) was added to an equal volume and samples were agitated for 5 min. Phases were separated by 30 min centrifugation (12,000g at 4 °C). Proteins were precipitated from the phenol phase (including those from PCI) by the addition of 5 volumes of 0.1 M ammonium acetate in methanol. Precipitated proteins were collected by centrifugation for 30 min (12,000g at 4 °C), washed twice with ammonium acetate in methanol and resuspended in resuspension buffer (3% SDS, 62.3 mM Tris-HCl, pH 8, 10% glycerol).

### RNA and protein blot analysis

For high molecular weight RNA analysis, 5-10µg of total RNA was separated on a 1.2% agarose MOPS-buffered gel with 2.2 M formaldehyde. RNA was partially hydrolyzed on gel with 5x gel volumes of 0.05N NaOH for 20 min. Gel was washed twice for 20 min. with 20X SSC, transferred overnight by capillarity to a HyBond-NX membrane (#RPN303, GE Healthcare) and UV-crosslinked for fixation. For high molecular weight RNA analysis by PAGE, 1-40µg of RNA (total, poly(A)+ or poly(A)-) were separated on a denaturing 4% polyacrylamide-urea gel, transferred to a HyBond-NX membrane by electroblotting and UV-crosslinked. For low molecular weight RNA analysis, 10-40µg of total RNA was separated on a denaturing 17.5% polyacrylamide-urea gel, transferred to a HyBond-NX membrane by electroblotting and chemically crosslinked (Pall and Hamilton 2008). Probes from PCR products were radiolabeled using the Prime-a-Gene kit (#U1100, Promega) in the presence of [α-^32^P]-dCTP (Hartmann Analytic) and oligo probes were radiolabeled by incubation of PNK (#EK0031, Thermo) in the presence of [α-^32^P]-ATP. Membranes were hybridized with these probes in PerfectHyb hybridization buffer (#H7033, Sigma) and detected on a Typhoon FLA 9500 (GE Healthcare) laser scanner. Oligonucleotides used for probe generation are listed in (Supplementary table 1).

Proteins were separated on SDS-polyacrylamide gels, transferred to Immobilon-P PVDF membranes (#IPVH00010, Millipore) by electroblotting and incubated with antibodies in 1X PBS with 0.1% Tween-20 and 5% nonfat dried milk. After incubation with HRP-conjugated secondary goat antibody against rabbit or rat primary antibodies (Sigma), detection was performed with the Clarity Max Western ECL substrate (#1705062, BIO-RAD) on a ChemiDoc Touch imaging system (BIO-RAD). Affinity-purified antibodies were used at the specified dilutions: GAG (1:2’000 (Oberlin et al. 2017)), GFP (1:5’000 Chromotek #3H9-100), GUS (1:1’000 Sigma Aldrich #G5545), H3 (1:10000 Abcam #ab1791), UGPase (1:2000 Agrisera #AS5 086). Protein loading was confirmed by Coomassie staining of membranes.

### Quantitative PCR

RNA was treated with DNaseI (#EN0521, Thermo Scientific) and cDNA was subsequently synthesized with the Maxima First-Strand cDNA Synthesis Kit (#K1641, Thermo Scientific), or RevertAid cDNA Synthesis Kit with Oligo(dT) (#K1612, Thermo Scientific). qPCRs were run on a LightCycler480 II (Roche) or a QuantStudio5 (Applied Biosystems) machine with the SYBR FAST qPCR Kit (KAPA Biosystems). Ct values were determined by the 2nd derivative max method of minimally two technical replicates for each biological replicate. Relative expression values were computed as ratios of Ct values between targets of interest and *ACT2* and/or *GAPC* reference mRNA unless otherwise indicated. *EVD* copy numbers were determined by direct qPCR on genomic DNA, comparing relative *EVD* and *ACT2* levels, normalized by their inherent copy numbers of two and one in WT plants, respectively. Oligonucleotides used are listed in Supplementary table 1.

### Separation of polyadenylated mRNA

Isolation of poly(A)+ from non-poly(A) RNA was performed from 75 ug of Trizol-extracted total RNA from floral buds, using the DynabeadsTM mRNA Purification Kit (Ambion Cat#.61006) following the manufacturer’s instructions. Non-polyA RNA was precipitated from the DynabeadsTM-unbound fraction and resuspended in the same volume (200 uL) as the poly(A)+ RNA fraction. Efficiency of the separation was confirmed by running aliquots of each fraction on a 1% agarose gel to monitor efficient depletion of rRNA in poly(A)+ fractions before downstream analysis.

### Cloning and mapping of 5’OH-ends

Non-canonical cleavage sites in *EVADE* transcript were mapped by a modified 5’ RACE method. Total RNA isolated from a pool of 2-3 weeks old plants extracted by standard protocols (See nucleic acid extraction section) were taken for RNA ligations after DNase I treatment ((#EN0521, Thermo scientific). RNA adapters with a 5’ inverted dT modification (See Supplementary Table 1) were ligated to the DNase-treated RNA by T4 RNA ligase 1 (#M0204S, New England Biolabs) to render the canonical cleavage products not available for subsequent ligation reaction. To map the cleavage products with a 5’ hydroxyl group, the RNA was subsequently ligated to an RNA adapter with a 3’ phosphate group by *RtcB* ligase (#M0458S, New England Biolabs). The ligated RNA was converted to cDNA with RevertAid first strand cDNA synthesis kit (#K1612, Thermo scientific) and a primer specific to the EVADE transcript (Supplementary Table 1). The cDNA was amplified by nested PCR by using primers from the adapter RNA and primers located ∼100 nucleotides downstream of each stalling site (all adaptor and primers sequences can be found in Suppl. Table 1). The PCR products were separated on an agarose gel and the DNA fragments were extracted from the gel by GeneJET gel extraction kit (#K0691, ThermoFisher scientific). The DNA fragments were cloned in pJET1.2 vectors by using CloneJET PCR cloning kit (#K1232, ThermoFisher scientific) and ∼50 colonies were screened for each potential cleavage site by Sanger sequencing technology.

### Small RNA sequencing

Small RNA sequencing of *35S:EVDwt* and *35S:GFP*-*EVD_int/ter_*-*GUS* was performed as follows. Total RNA was resolved on a 17.5% polyacrylamide-urea gel and sizes between 18 - 30 nt were excised, eluted overnight in elution buffer (20mM Tris–HCl (pH 7.9), 1 mM EDTA, 400 mM ammonium acetate, 0.5% (w/v) SDS) and collected by precipitation with equal volumes of isopropanol. RNA was quantified using the Qubit™ RNA HS Assay Kit (Thermo Scientific) and subsequently cloned using the Small RNA-Seq Library Prep Kit (Lexogen). Sequencing was performed on an Illumina HiSeq 4000 machine.

### RIBO-seq

For RIBO-seq libraries frozen inflorescence tissue was ground in digestion buffer (100 mM Tris·HCl (pH 8), 40 mM KCl, 20 mM MgCl*_2_*, 2% (v/v) polyoxyethylene (10) tridecyl ether, 1% (v/v) de-oxycholic acid, 1 mM DTT, 10 unit/mL DNase I (Thermo Scientific), 100 μg/mL cycloheximide). Precleared solutions were incubated with 650 U RNase I (Ambion) for 45 min at 25°C. Nuclease digestion was stopped by the addition of 10 μl SUPERase In RNase Inhibitor (Ambion). Resulting monosomes were purified by ultracentrifugation of the lysate on a sucrose cushion (1 M sucrose, 20 mM HEPES (pH 7.6), 100 mM KCl, 5 mM MgCl2, 10 μg/ml cycloheximide, 10 units/ml RiboLock (Thermo Scientific) and cOmplete protease inhibitor cocktail (Roche) for 4 hours at 250’000 g in 4°C. RNA was extracted using the TRIzol RNA extraction described above and treated with 10 U PNK (Thermo Scientific) for 30 min. Ribosomal RNA depletion was performed using the RiboMinus Plant Kit (Thermo Scientific) and libraries were generated as above, except that the 25 - 32 nt RNA fraction was excised from the denaturing polyacrylamide gel prior to RNA ligation.

### nanoPARE

NanoPARE library preparation and analysis was performed following the protocol from Schon et al. 2018 (Schon *et al*, 2018). Briefly, 10ng of total RNA was isolated from inflorescences. Two biological replicates each of Col-0 and *ddm1-2* were used for reverse transcription. After 9 cycles of PCR pre-amplification, 5ng aliquots of cDNA were separately tagmented and amplified using either standard Smart-seq2 Tn5 primers or 5’-end enrichment primers. The resulting Smart-seq2 and nanoPARE libraries were sequenced on an Illumina HiSeq 2500 using paired-end 50bp reads and single-end 50bp reads, respectively.

### Seed counting

Plants germinated and grown in parallel under the same conditions were individually covered with paper bags before the maturation of siliques and harvested upon ripening. Total amount of seeds from each plant was counted twice with a C3 High Sensitive Seed Counter (Elmor).

### Data Analysis

Analysis of sRNA sequencing is based on the following workflow. Reads were trimmed using bbduk (BBTools: <u>sourceforge.net/projects/bbmap/</u>, version 38.41; ktrim=r k=23 mink=11 hdist=1) mapped against the TAIR10 Arabidopsis genome with STAR (Dobin *et al*, 2013) (version 2.5.2a; --outFilterMismatchNoverLmax 0.05 –outFilterMatchNmin 15 --outFilterScoreMinOverLread 0 --alignIntronMax 500 --alignIntronMin 50 -- outFilterMultimapNmax 50), quantified using Rsubread (Liao *et al*, 2013) (version 1.20.6; allowMultiOverlap=T,largestOverlap= T, isPairedEnd=F, strandSpecific=1, countMultiMappingReads=T, fraction = T) and differential analysis using DESeq2(Love *et al*, 2014) (version 1.10.1). Reads were split in different lengths with Samtools (Li *et al*, 2009) (version 0.1.19) and locus coverage amongst those read length was visualized using BEDtools (Quinlan & Hall, 2010) (version 2.15.0) and R cran (version 3.2.5).

RIBO-seq libraries were analyzed as follows. Reads were trimmed of adapter sequences with bbduk as above. Reads mapping to rRNA loci using Bowtie2 (Langmead & Salzberg, 2012) (version 2.2.1; -k 1 -x) were discarded from further analysis. Subsequent mapping and quantification were performed as for the sRNA sequencing analysis using STAR (Dobin *et al*, 2013) and Rsubread (Liao *et al*, 2013) as above, but reads were mapped to both *Arabidopsis* genome and transcriptome sequences. Quality control of the RIBO-seq libraries was performed with the riboWaltz (Lauria *et al*, 2018) (version 1.1.0) package. P-site occupancies were estimated using the RiboProfiling (Popa *et al*, 2016) (version 1.0.3) package based on 5’ read offsets determined by the coverage profile around start codons dependent on read lengths. Codon occupancies were compiled for all three possible frames to generate a single codon occupancy score. A ribosomal stalling score at each codon position was defined as the ratio of observed over expected counts, where the expectation was the mean of occupancy counts over the entire transcript. To improve quality of the assessment, only the most translated isoform per gene and only isoforms with a minimal read coverage of 70% were considered. Codon dwell time was estimated as the mean value of log-normalized codon occupancies per individual transcript and codon usage was estimated from the subset of genes considered translated. Stop codons and stop codons containing di-codons were excluded from the analysis. Data were visualized using R cran and the packages Gviz (Hahne & Ivanek, 2016) and ggplot2 (Wickham).

### Data Access

Sequencing data generated in this study are accessible on the Gene Expression Omnibus (GEO) under the accession number <u>GSE167484</u>. Data from previous studies including sRNA sequencing in *ddm1* & *ddm1 rdr6*, *ddm1* & *ddm1 dcl1,* isoform specific sequencing data of total and polysome associated mRNA in TE de-repressed backgrounds are found under the accession numbers <u>GSE41755</u> (Nuthikattu *et al*, 2013), <u>GSE52952</u> (Creasey *et al*, 2014), <u>GSE93584</u> (Oberlin *et al*, 2017) and <u>PRJNA598331</u> (Kim *et al*, 2021). Every other raw data used in this study (including raw image files, qPCR data, Sanger sequencing traces,…) has been deposited in Zenodo (www.zenodo.org) under the DOI: <u>10.5281/zenodo.4751668.</u>

