## Supplementary Figures 1-11 and Supp_table for "Innate, translation-dependent silencing of an invasive transposon in Arabidopsis"

### Supplemental Information 1

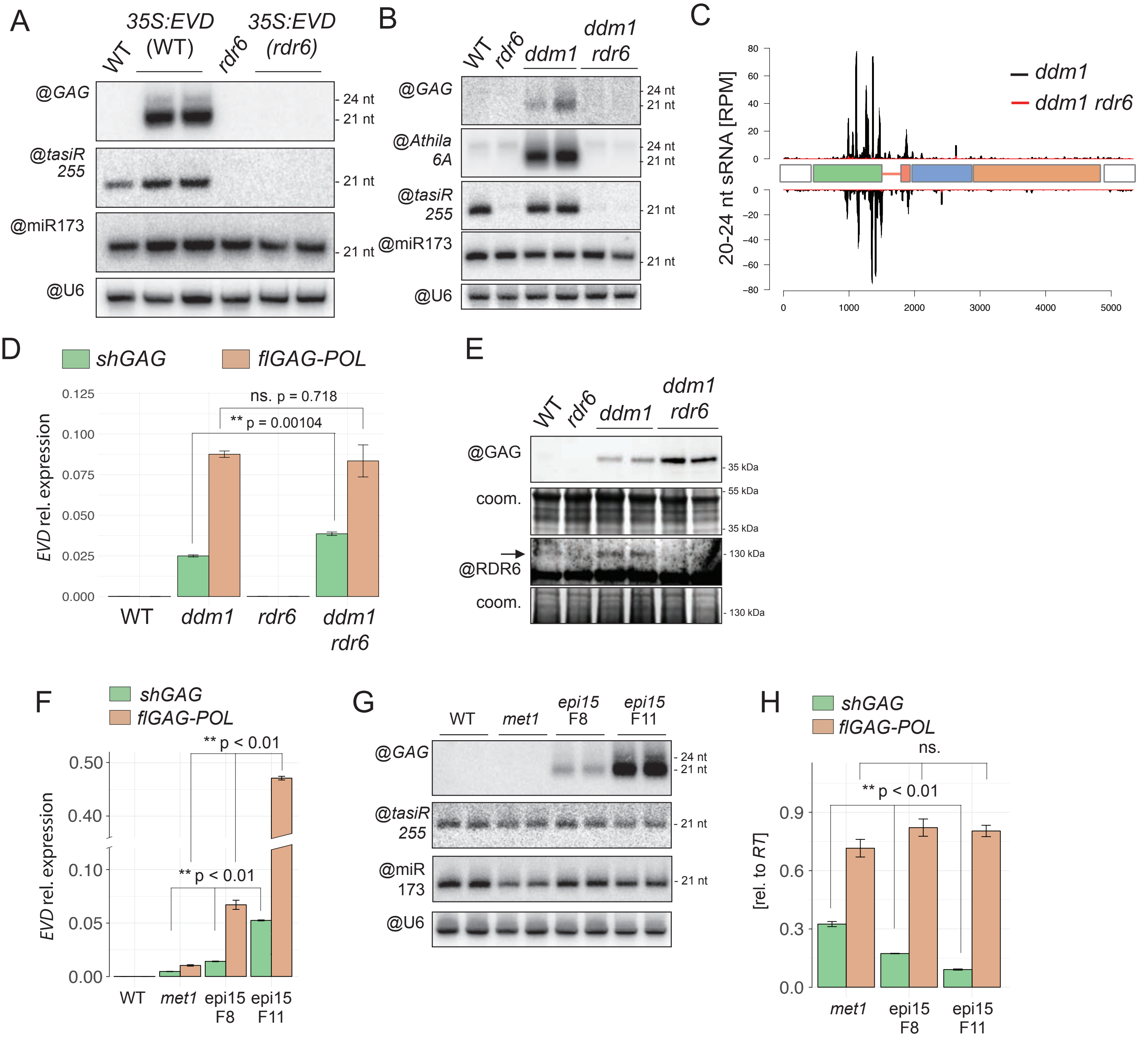


**Supplemental Figure 1. The EVD spliced and prematurely terminated shGAG mRNA is a trigger and target of RDR6-dependent silencing. (A)** Low molecular weight RNA analysis from *35S:EVDwt* in the WT or *rdr6* background with probes against *GAG* or tasiR255 as a control for the *rdr6* mutation or miR173 and U6 as loading controls. **(B-E)** Characterization of endogenous *EVD* in *ddm1* or *ddm1 rdr6* backgrounds. (B) Low molecular weight RNA analysis. *Athila 6A*, tasiR255, miR173 and U6 are used as controls. (C) 20-24 nt siRNA profile on the *EVD* locus. RPM: Reads per million. (D) Relative expression of *shGAG* versus *flGAGPOL* mRNAs. (E) Western analysis of GAG and RDR6. Coomassie (coom.) blue staining provides a loading control. **(F-H)** Endogenous *EVD* expression and *GAG* siRNA accumulation caused by increased copy number in WT, *met1* or *met1*-derived epiRIL *epi15* from generation F8 to F11. (F) qPCR analysis of the two mRNA isoforms relative to *ACT2*. (G) Low molecular weight RNA analysis, with controls as in (B). **(H)** qPCR analysis of the spliced and unspliced transcripts levels relative to *EVD RT* expression, reflecting the ratios to the full-length isoform. In all panels: qPCR was performed on three biological replicates and normalized to *ACT2* unless indicated. Error bars represent the standard error. (**) = p-value < 0.01 (two-sided t-test against corresponding controls).

### Supplemental Information 2

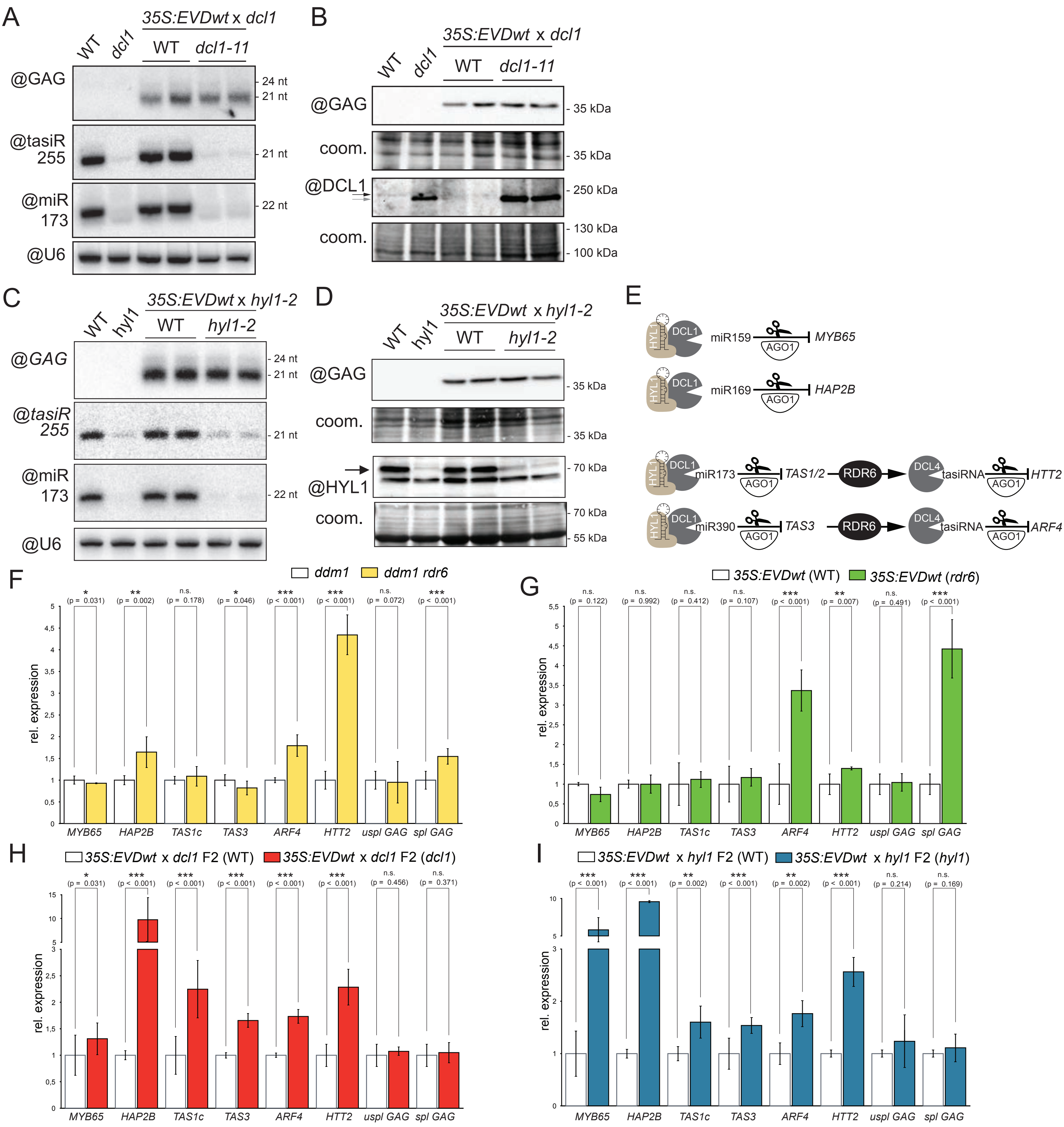


**Supplemental Figure 2. miRNA-independent silencing of *EVD* in *ddm1* and *35S:EVDwt* overexpression lines.** **(A-B)** Analysis of *35S:EVDwt* in the *dcl1-11* versus WT background. (A) Low molecular weight RNA analysis with probes against *GAG* or miR173, tasiR255 and U6 as controls. (B) Western analysis of GAG and DCL1 . Note that WT DCL1 (black arrow) is slightly larger than the mutated dcl1-11 protein (gray arrow). dcl1-11 is also upregulated due to the loss of the miR162-mediated negative feedback loop controlling *DCL1* levels. The coomassie (coom.) blue-stained membrane is shown as a loading control. **(C-D)** Analysis of *35S:EVD* in *hyl1-2*. (C) Low molecular weight RNA analysis of *GAG* as well as miR173, tasiR255 and U6, used as controls. (Continues on next page)

(D) Western analysis of GAG and HYL1. The coomassie (coom.) blue-stained membrane provides a loading control. **(E)** Succinct schemes for DCL1- and HYL1-dependent regulation of miRNA targets *MYB65* and *HAP2B* (top) and for miRNA-RDR6-dependent initiation of tasiRNA biogenesis and regulation of targets *HTT2* and *ARF4* (bottom). **(F - I)** Relative expression levels of multiple miRNA targets, tasiRNA precursors, tasiRNA targets and EVD transcript in *35S:EVDwt* in the indicated genetic backgrounds as used in Fig.1, Fig.2, Supp.Fig.1 and Supp.Fig.2. In all panels: qPCR was performed in three biological replicates. Error bars represent the standard error. (ns.) = non-significant, (*) = p-value < 0.05 , (**) = p-value < 0.01, (***) = p-value < 0.001, (two-sided t-test against corresponding controls).

### Supplemental Information 3

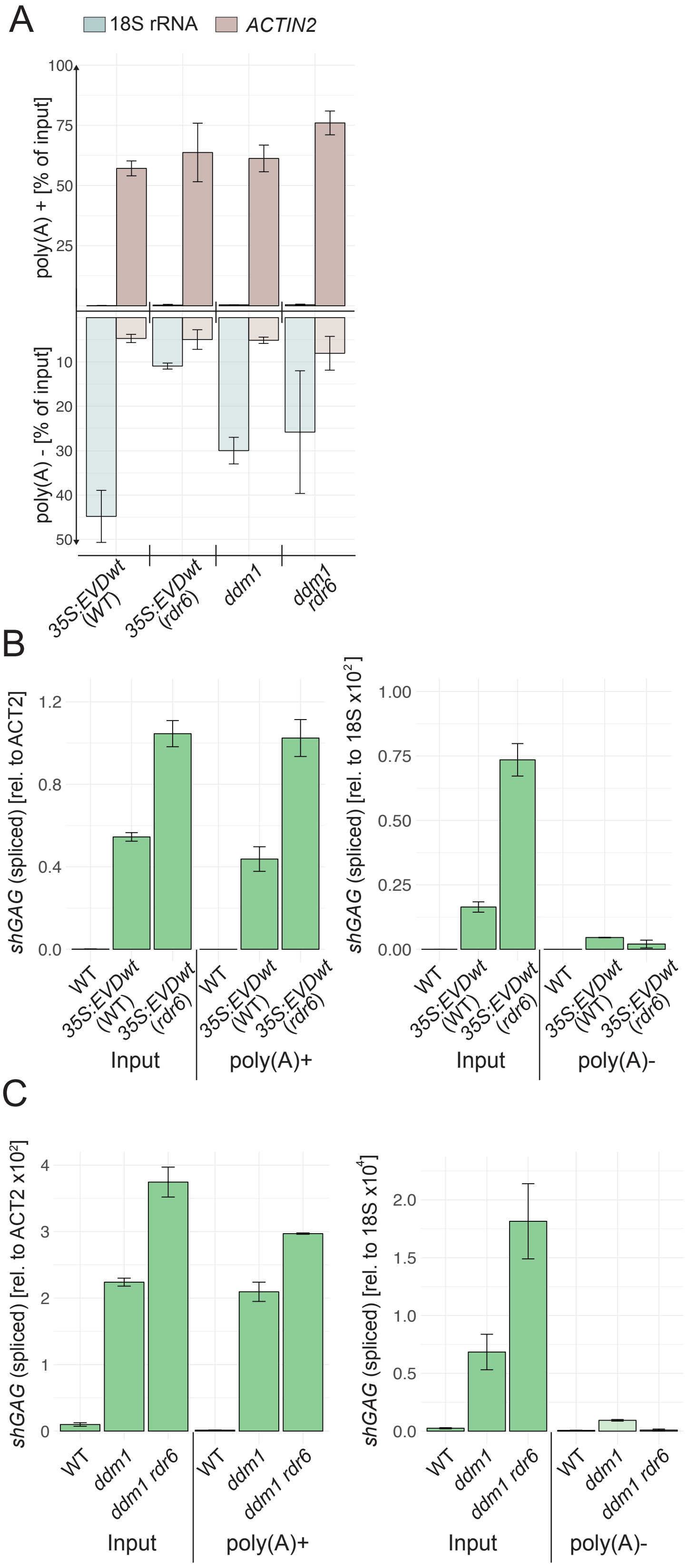


#### Defective shGAG poly(A)-tailing does not underlie siRNA production.

Transgene mRNAs lacking a poly-adenylation (poly(A)) tail as a result of aberrant transcription stimulate RDR6 activity in vivo. To test if potential poly(A) defects could explain RDR6 affinity for the spliced shGAG mRNA, total polyadenylated (poly (A)+) versus non-polyadenylated (poly(A)-) RNA was fractionated from *35S:EVDwt* tissues in either the WT or *rdr6* background. This was confirmed in *ddm1* and *ddm1 rdr6* non-transgenic plants in which EVD is epigenetically reactivated (Fig.1, S1). In both settings, *shGAG* was near-exclusively poly(A)+ (Fig.S3A-C). Accordingly, the increased *shGAG* mRNA levels in rdr6 were contributed by the global poly(A)+, not poly(A)-, fraction (Fig.S3B,C). Therefore, aberrant transcription leading to poly(A)-tail-deficiency is unlikely to stimulate RDR6 recruitment specifically on *shGAG*.

**Supplemental Figure 3. Effect of rdr6 on endogenous and transgenic polyadenylated versus non-polyadenylated EVD RNA levels. (A)** qPCR assessment of poly(A)- versus poly(A)+ RNA fractionation after two subsequent oligo(dT) purifications. Quantification involves 18S rRNA (poly(A)-) and *ACT2* (poly(A)+) controls relative to input (total RNA). Note that two poly(A) separation steps are not expected to yield a 100% recovery. **(B)** *shGAG* mRNA quantification in input, poly(A) + and poly(A) - fractions relative to *ACT2* (poly(A)+) and 18S rRNA (poly(A)-) in *35S:EVDwt* in the WT or *rdr6* background. **(C)** Same as in (B) but in the *ddm1* versus *ddm1 rdr6* background. qPCR was performed in biological duplicates and error bars represent the standard error.

### Supplemental Information 4

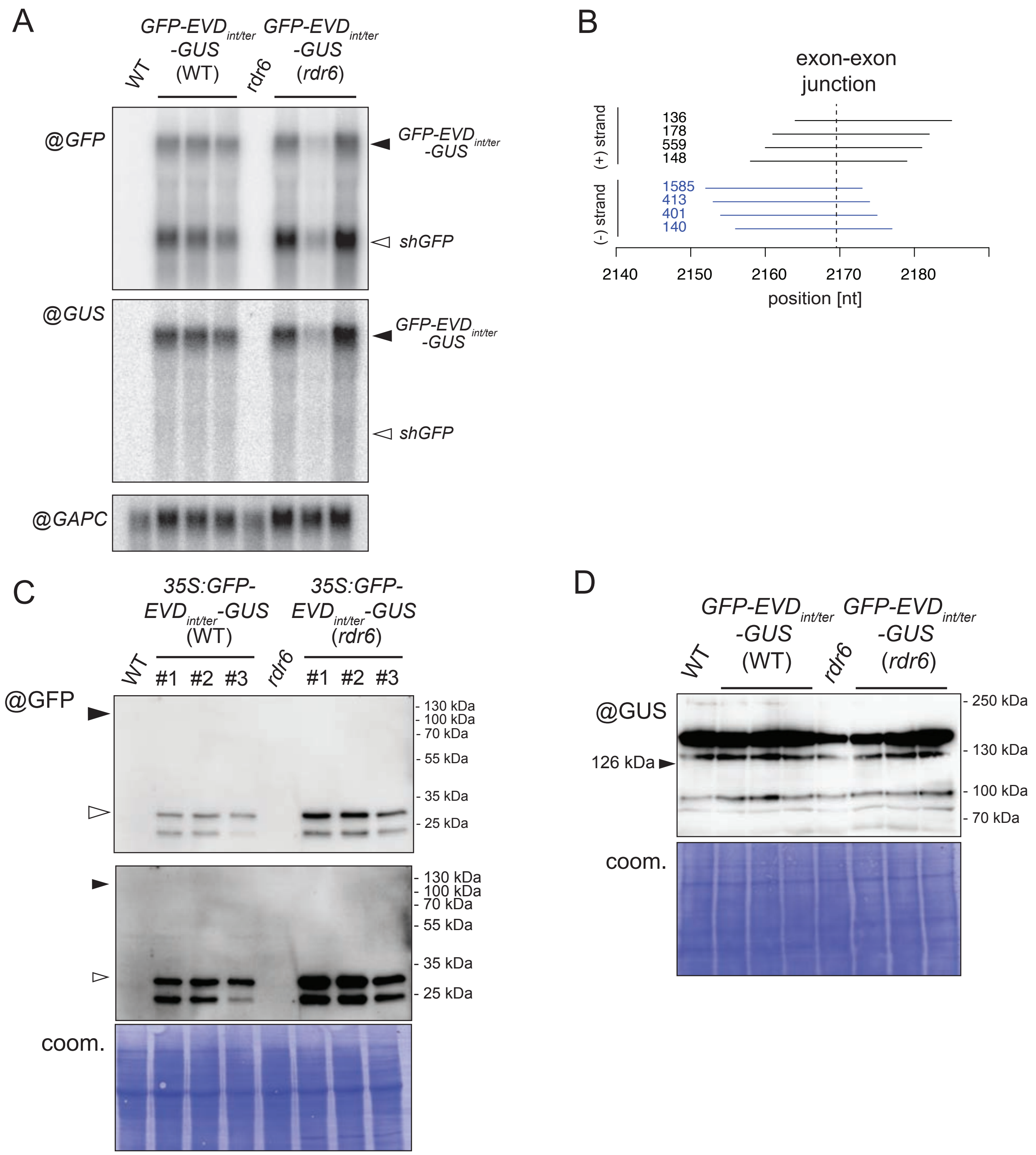


**Supplemental Figure 4. Molecular characterization of 35S:GFP-EVDint/ter-GUS expressed in the WT or rdr6 background. (A)** High molecular weight RNA analysis probing for GFP and GUS regions in three independent transgenic lines in either WT or rdr6 alongside with non-transformed controls. GAPC probing is used as a loading control. **(B)** sRNA reads mapping at the splice junction of the EVD intron on the positive (+) and negative (-) strand alongside their abundance in the WT background. No reads covering the intron junction were found in rdr6. **(C-D)** western analysis of GFP (C) and GUS (D). GFP-only protein (white arrow; ~30 kDa), but not the GFP-GUS fusion protein (back arrow; 126kDa) was detected. Coomassie staining (coom.) of the membrane serves as loading control.

### Supplemental Information 5

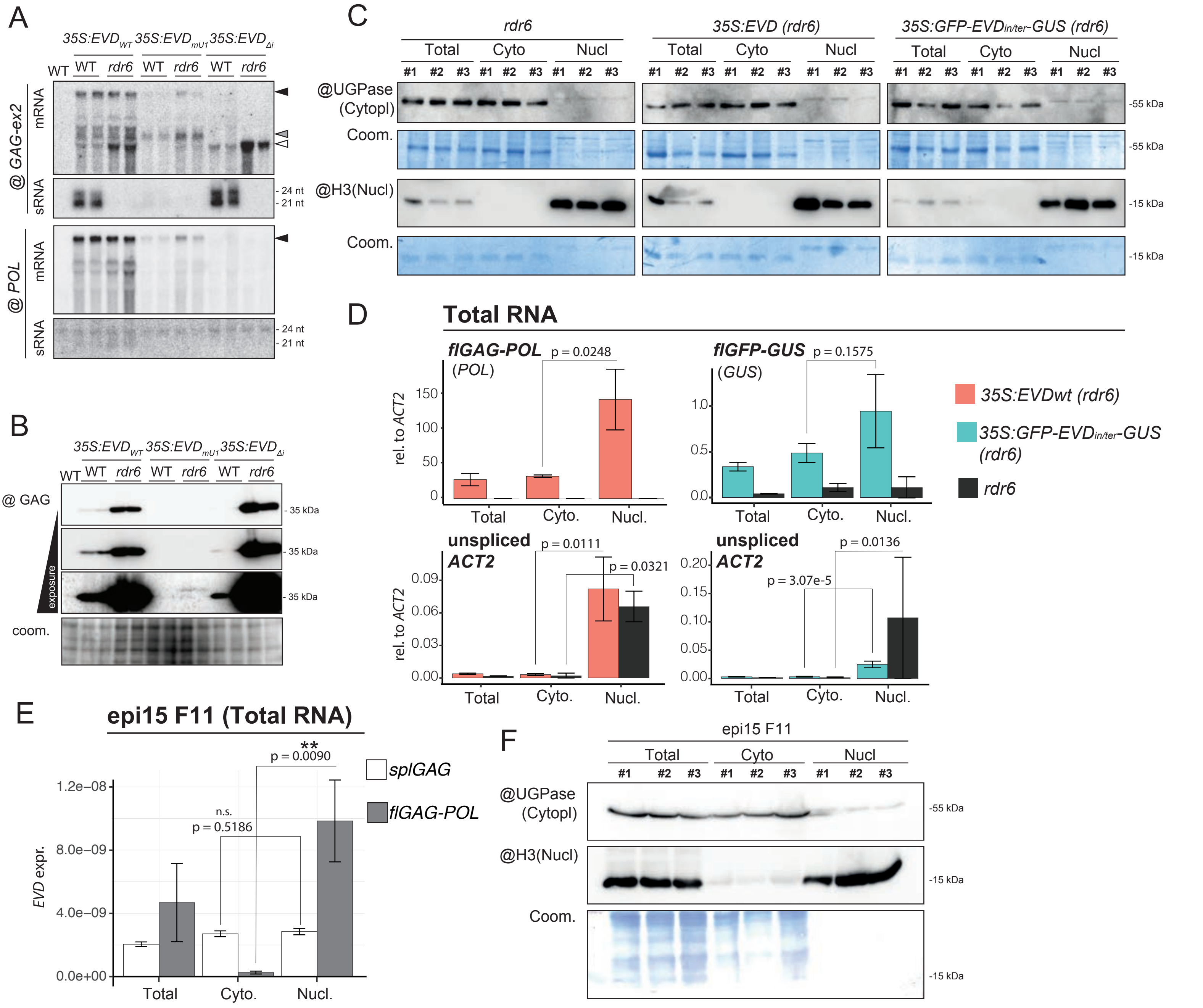


**Supplemental Figure 5. Impact of splicing and premature termination on EVD silencing and mRNA localization.** **(A)** High and low molecular RNA analysis of the *shGAG* (GAG-ex2 probe) or *flGAG-POL* (POL probe) transcripts. The mRNA isoforms are labelled with arrows on the side or with an asterisk and correspond to the transcripts depicted in Fig.3A-C. Loading controls are displayed in Fig.3E. **(B)** Same western analysis of the GAG protein as presented in Fig.3F, but with three increasingly higher exposure times and Coomassie loading control (coom.). **(C)** Quality assessment of nucleo-cytosolic fractionations from *rdr6*, *35S:EVD(rdr6)* and *35S:GFP-EVDint/ter-GUS (rdr6)* by western analysis. UGPase and HISTONE 3 (H3) were used as cytoplasmic and nuclear protein markers, respectively, in protein extracted from total, cytoplasmic (Cyto) or nuclear (Nucl) fractions from three independent experiments. Coomassie staining (coom.) of the membranes serves as loading control. **(D)** Nucleo-cytosolic distribution of unspliced *ACT2*, *35S:EVD* and *35S:GFP-EVDint/ter-GUS* full-length RNA in the rdr6 background relative to that of spliced *ACT2* by qPCR. RNA extracted from Total, nuclear and cytoplasmic fractions was reverse transcribed with oligo(dT) to account only for polyadenylated RNA. **(E)** Nucleo-cytosolic distribution of endogenous *EVD* RNA isoforms in the *epi15* F11 analyzed by qPCR from total, cytoplasmic and nuclear fractions. **(F)** Quality assessment of nucleo-cytosolic fractionations, as in C, from epi15 F11 in the three biological replicates used in E. In D and E qPCR was performed on n=3 biological replicates; bars: standard error. (*) = p-value < 0.05; (**) = p-value < 0.01; n.s = non significant (p-value > 0.05) (two-sided t-test between indicated samples).

### Supplemental Information 6

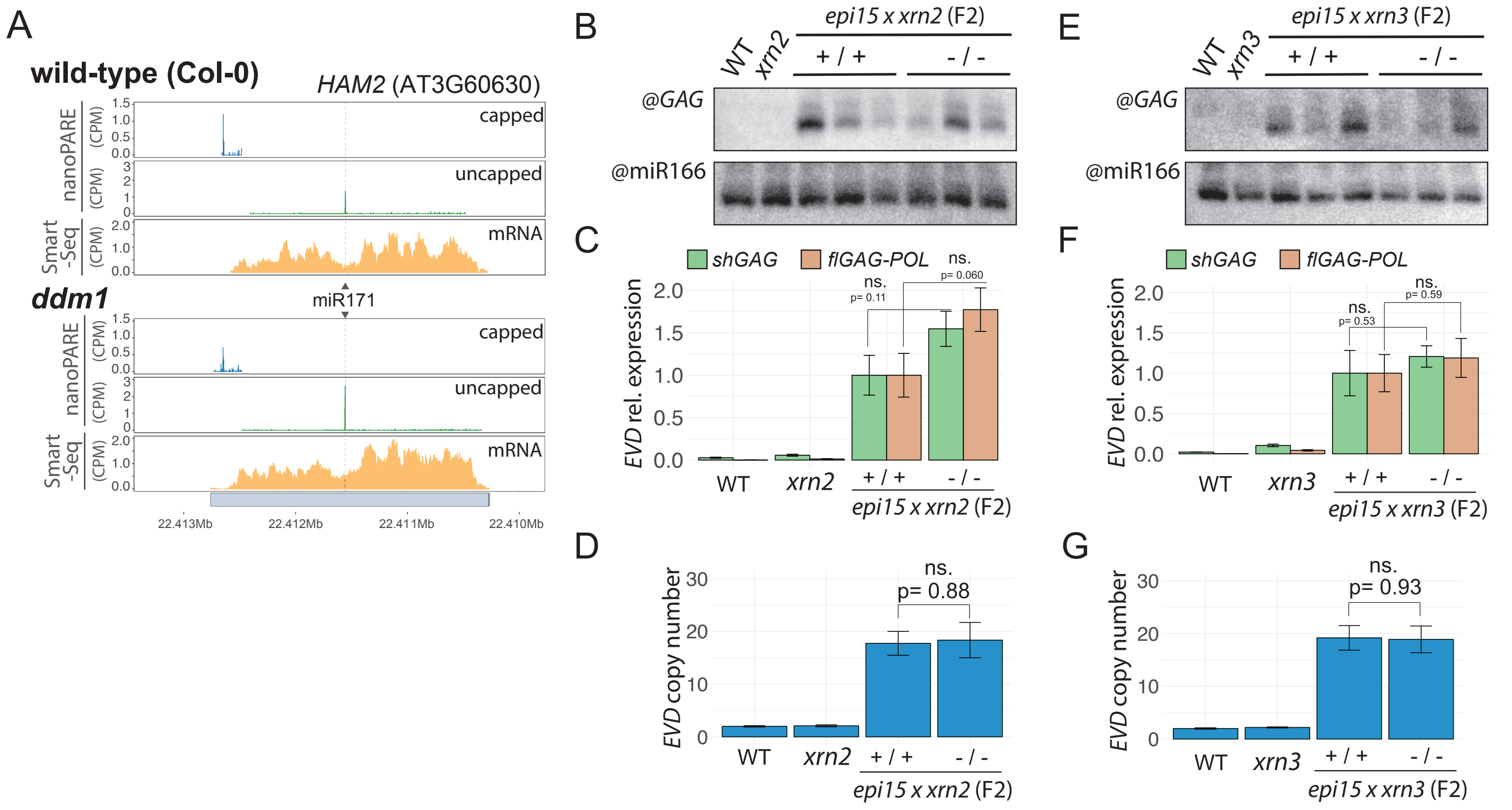


**Supplemental Figure 6. Neither *xrn2* nor *xrn3* alter *EVD* siRNA production, *EVD* expression or *EVD* copy number.** **(A)** Capped and uncapped 5'ends mapping with nanoPARE on the miR171 target *HAM2* in WT and *ddm1*, along with the corresponding Smart-seq coverage (mRNA). Capped and uncapped defined respectively as reads overlapping, or not, with capped 5' clusters (reads containing 5' untemplated G) **(B-D)** Impact of loss-of-XRN2 function on active *EVD* assessed in homozygous *xrn2* and WT backgrounds in F2 plants from a cross between *xrn2* and *epi15* plants undergoing active EVD mobilization. (B) Low molecular weight RNA analysis, with miR166 as a loading control. (C) Relative expression levels of *shGAG* and *flGAGPOL*. (D) EVD genomic copy number determined by qPCR assay on genomic DNA. **(E-G)** Same as (B-D) but with the *xrn3* mutation. In all panels, qPCR was performed on three biological replicates for controls and three independent WT and mutant F2 lines. Error bars display standard errors. (ns.) = non-significant, (*) = p-value < 0.05, (**) = p-value < 0.01 (two-sided t-test between indicated samples).

### Supplemental Information 7

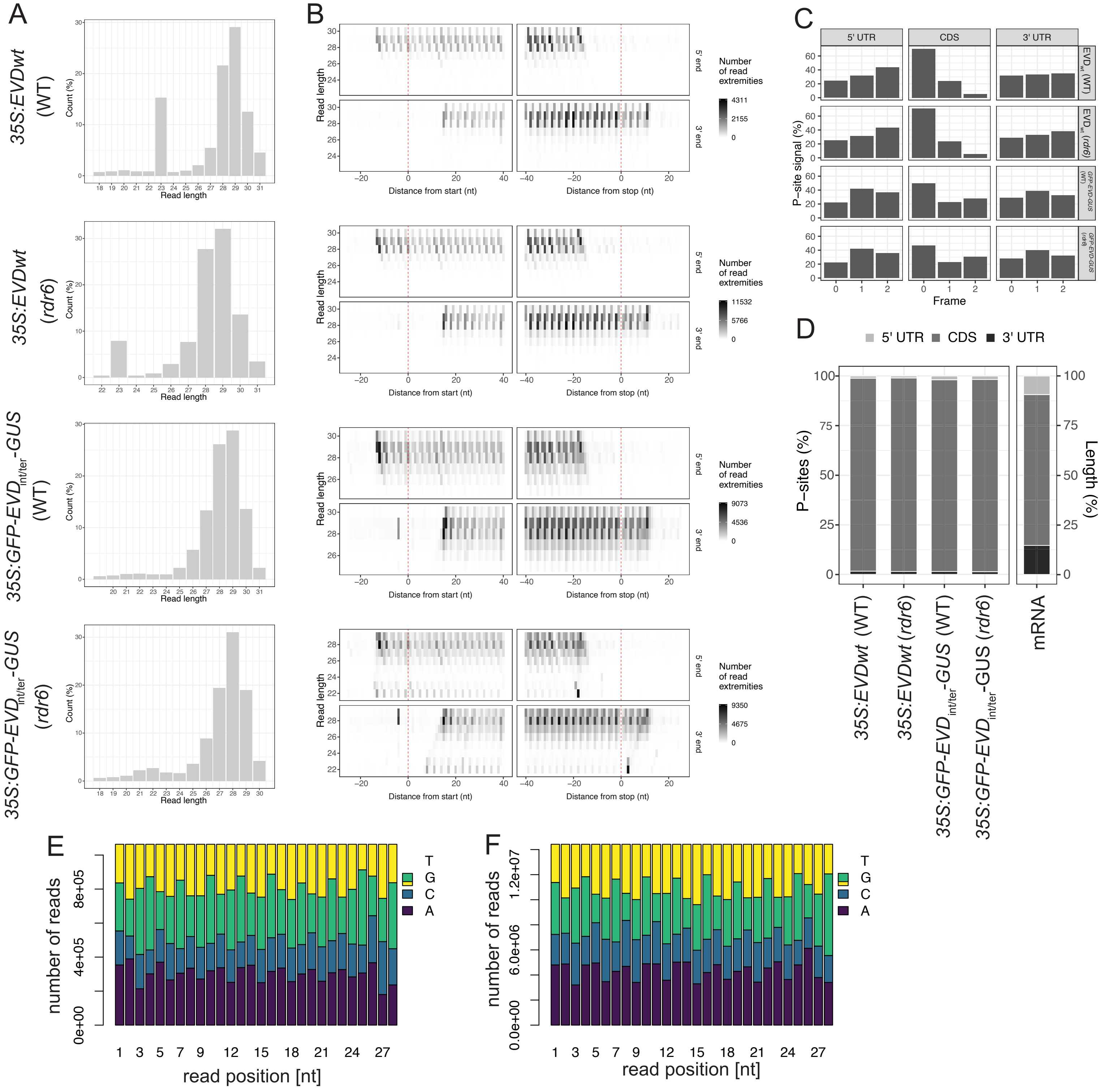

**Supplemental Figure 7. Quality control of the ribosomal footprinting libraries in the indicated genetic backgrounds. (A)** Read-length profiles of reads mapping to transcripts of protein coding genes in each of the indicated library. **(B)** Footprinting periodicity and offset across read lengths of 24 to 30 nt for each library. The translational start (left) and stop (right) sites are indicated with red vertical lines. Reads densities on the 5′ end (top) and the 3′ end (bottom) are displayed. **(C)** Triplet periodicity was captured inside coding sequences (CDS) but no in untranslated regions (UTRs) upon assigning reads counts to the three positions within each codon. **(D)** Significant enrichment of reads coverage from CDS as opposed to both 5′ and 3′ UTRs compared with the average length of those elements within Arabidopsis transcripts. **(E)** Nucleotide composition at each position of 28 nt long reads from the 35S:EVDwt library (WT background) used in this work. **(F)** Same as in (E) but with a library prepared according to the original RIBO-seq protocol described by Ingolia et al. 2009, presented here for comparison.

### Supplemental Information 8

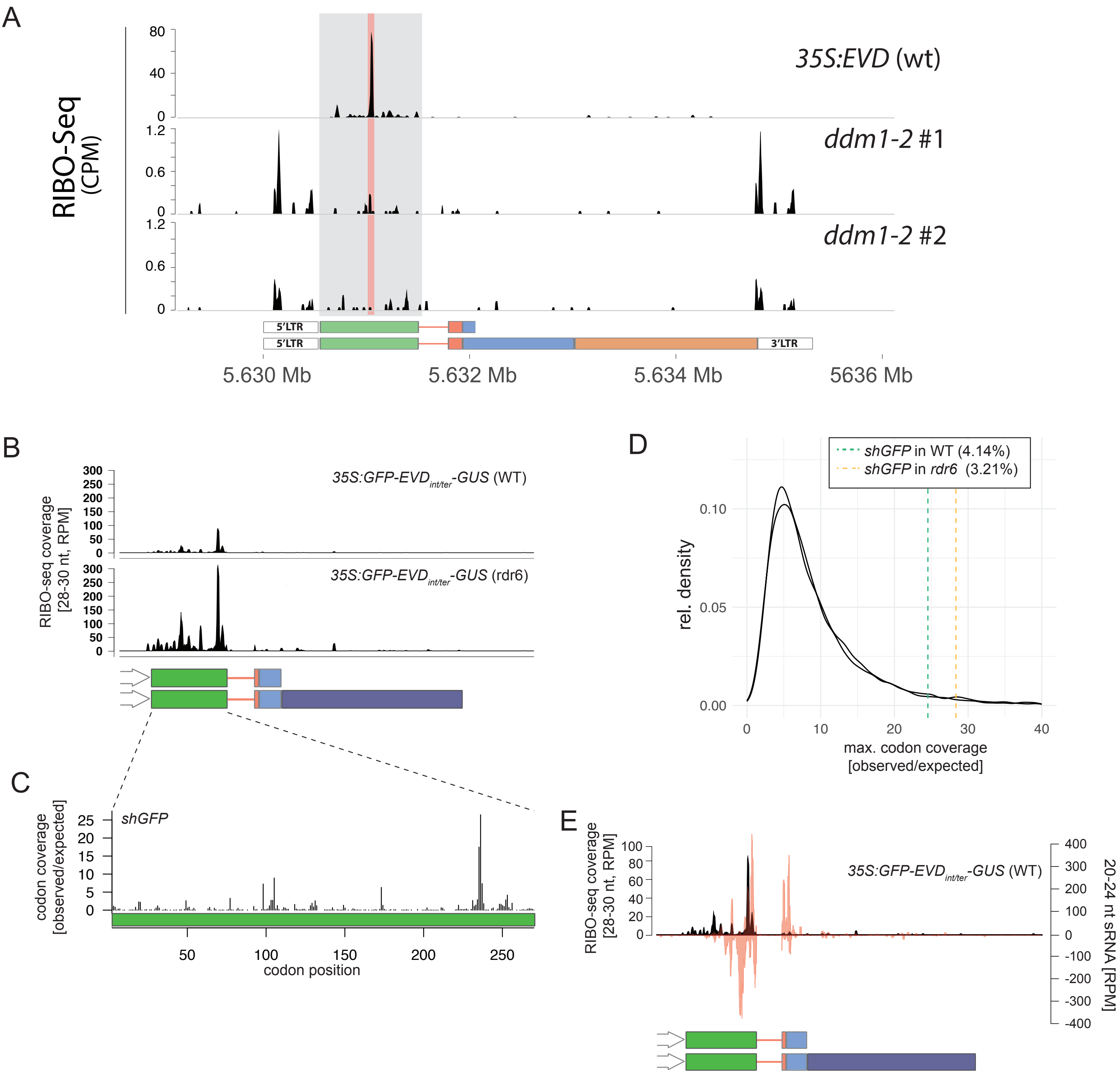


**Supplemental Figure 8. Ribosome footprints on 35S:GFP-EVDint/ter-GUS. (A)** *EVD* RIBO-seq coverage profiles on the *35S:EVD* in WT and two *ddm1-2* libraries. The *GAG* coding region is highlighted in grey, and the conserved stalling site in both *35S:EVD* in WT and *rdr6* (Fig.7A) is highlighted in red. CPM: Counts per million. **(B)** RIBO-seq coverage profiles on the 35S:GFP-EVDint/ter-GUS in WT and rdr6. RPM: Reads per million. **(C)** Ribosomal footprints compiled to display codon occupancy at the P-sites over the GFP sequence. Observed coverage at each codon position was divided by the expected mean coverage along the entire GFP coding sequence. **(D)** Maximal individual codon coverage over the expected coverage for all translated transcripts of Arabidopsis. Vertical lines indicate the strength of stalling sites on shGFP in the WT or rdr6 background. Percentages specify the proportion of transcripts with more pronounced stalling events than the shGFP ones. **(E)** Overlay between 35S:GFP-EVDint/ter-GUS siRNAs in WT and RIBO-seq profiles in the *rdr6* background.

### Supplemental Information 9

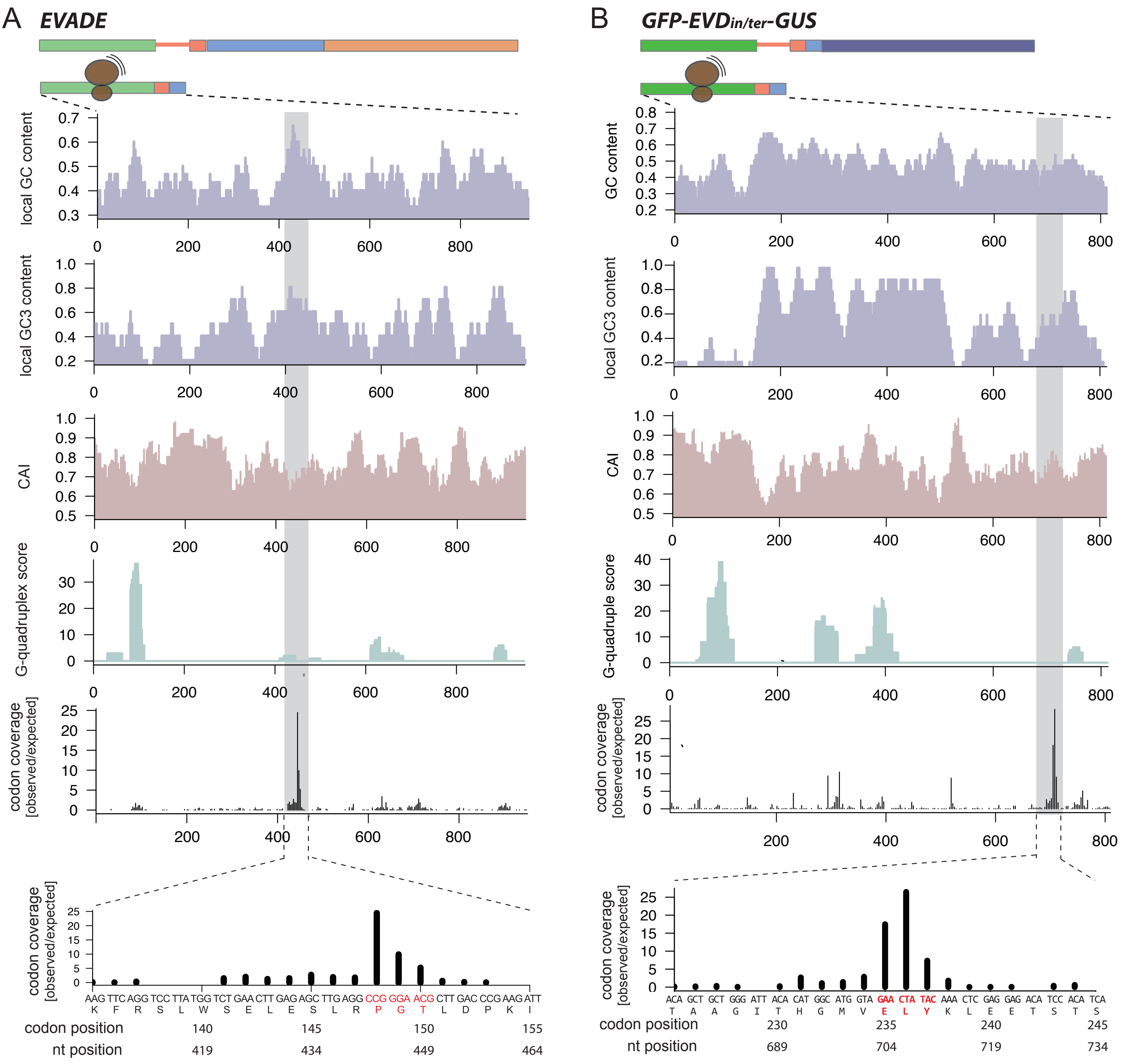


**Supplemental Figure 10. Sequence properties with the potential to impact translation of shGAG (A) and shGFP (B) transcripts.** For local GC content, GC content at codon position three and codon adaptivity index, sequences were analysed as 30 nucleotide long sliding windows shifting by three nucleotide steps representing one codon shifts. G-quadruplex scores were predicted as previously reported with pqsfinder. Codon coverage represents the ribosomal footprints compiled to display codon occupancy at the P-sites over the *shGAG* and *shGFP* sequence. Observed coverage at each codon position was divided by the expected mean coverage along the entire GFP coding sequence.

### Supplemental Information 10

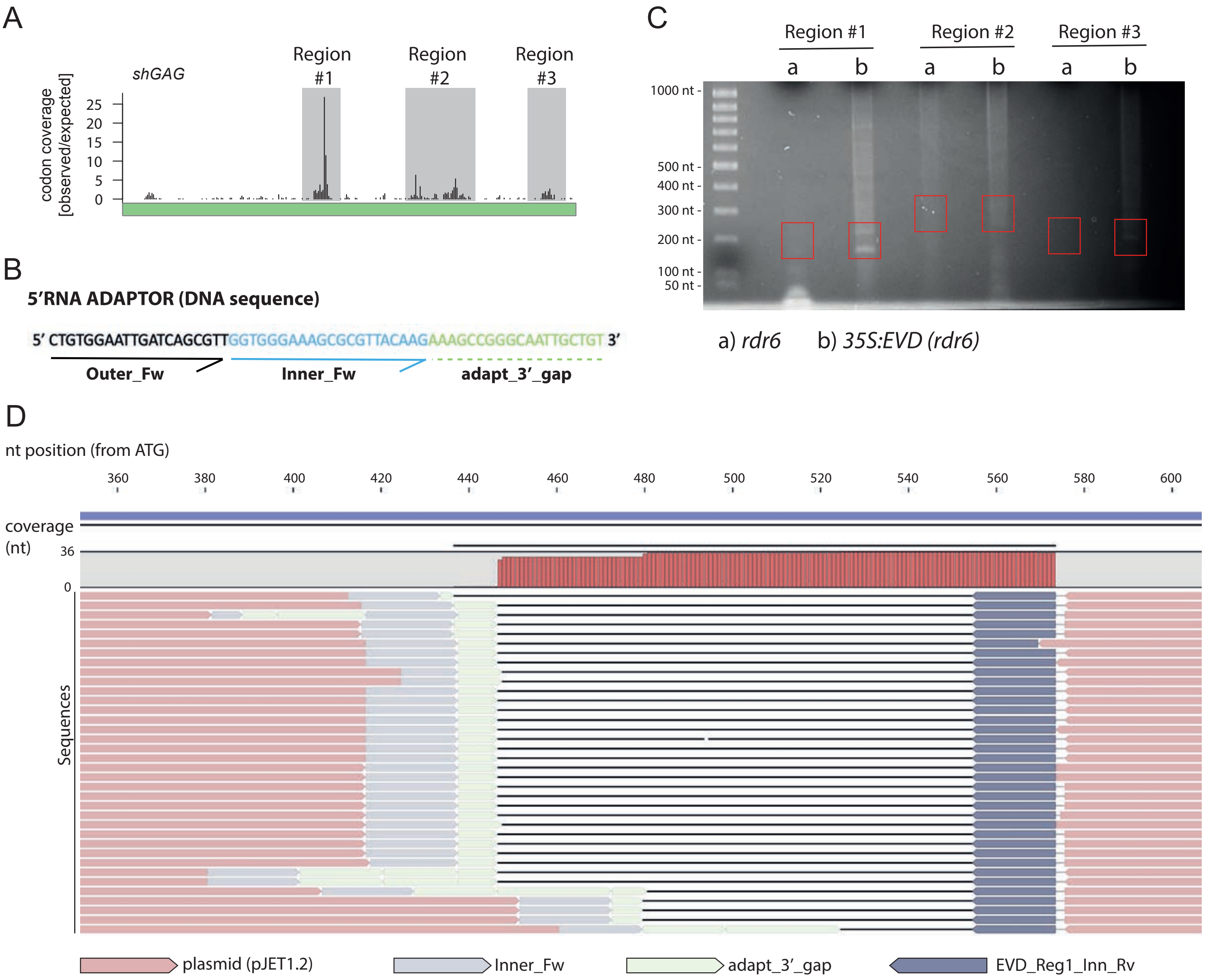


**Supplemental Figure 9. Cloning and Mapping of *EVD* 5'OH-ends.**
**(A)** Ribosome footprints over *EVD shGAG* transcript. The three regions inspected for atypical 5'OH ends are higlighted in grey. **(B)** Sequence of the RNA adaptor used to ligate to 5'OH ends with *RbtC*. Binding sites for outer and inner primers for nested PCR are indicated. **(C)** Ethidium Bromide staining of amplification products from second nested PCR in *rdr6* and *35S:EVD (rdr6)* resolved on an agarose gel. Independently of the presence/absence of bands, for all PCRs, gel was excised (red squares) for DNA extraction and cloning at the expected size for any potential amplicon within the regions highlighted in A. **(D)** Mapping, alignment and annotation of positive clones/colonies. To discard potential PCR artifacts, only clones displaying at least 1/4 of adaptor sequence between adaptor Inner_Fw primer and *EVD* sequence (adapt_3' _gap, B) were taken for the analysis. Nucleotide-resolution mapping of the 5'ends is shown in Fig.6F.

### Supplemental Information 11

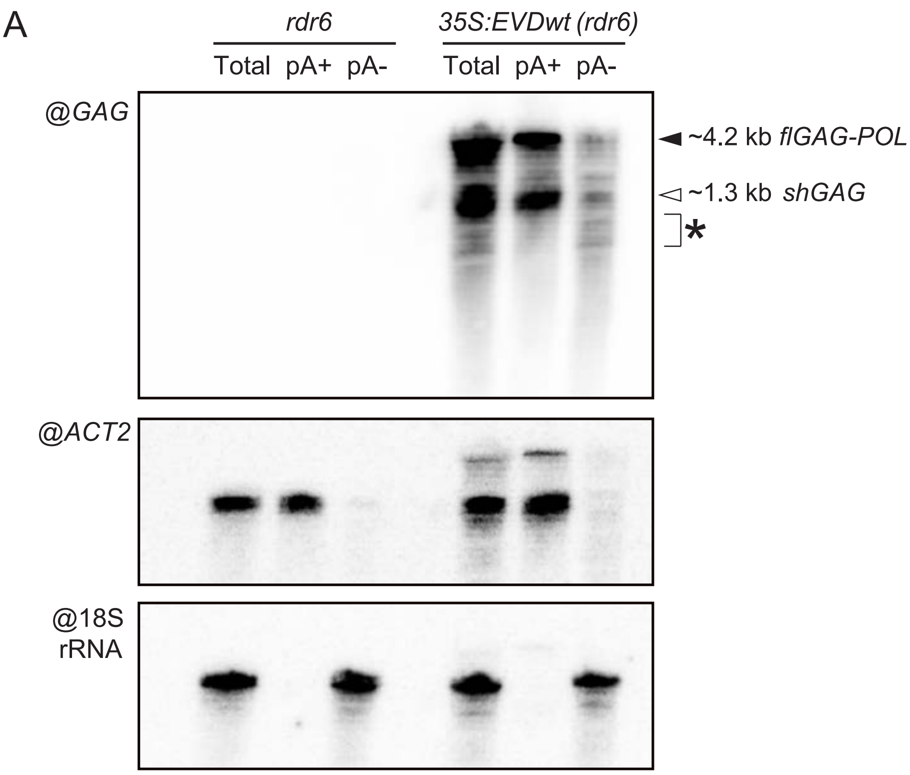


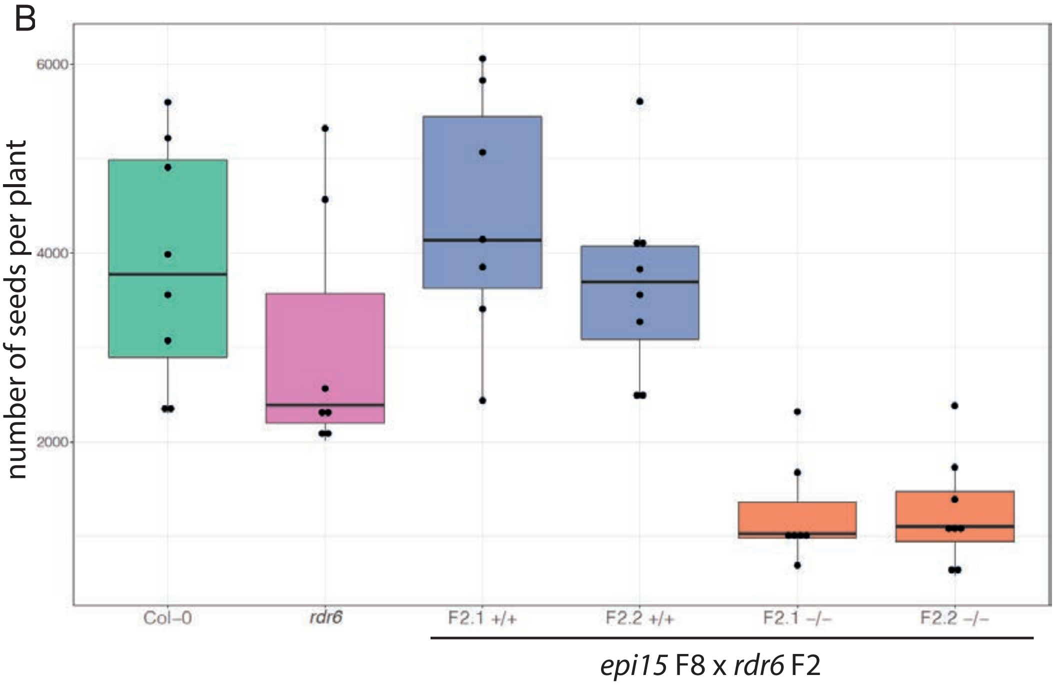


**PAIRWISE COMPARISON P-VALUES:**

| | Col-0 | F2.1 (-/-) | F2.1 (+/+) | F2.2 (-/-) | F2.2 (+/+) |
|---|---|---|---|---|---|
| F2.1 (-/-) | ** 0.0064 | —— | —— | —— | —— |
| F2.1 (+/+) | n.s. 1.0000 | ** 0.0064 | —— | —— | —— |
| F2.2 (-/-) | ** 0.0044 | n.s. 1.0000 | ** 0.0044 | —— | —— |
| F2.2 (+/+) | n.s. 1.0000 | ** 0.0044 | n.s. 1.0000 | ** 0.0023 | —— |
| *rdr6* | n.s. 1.0000 | * 0.0326 | n.s. 0.5099 | * 0.0196 | n.s. 1.0000 |

**Supplemental Figure 11. (A)** Separation of poly(A)+ and poly(A)- RNA from *35S:EVDwt* in rdr6 on a 4% PAGE gel hybridized with a probe against *GAG* to detect both *shGAG* and *flGAG-POL* mRNA isoforms. Putative stalling-linked RNA cleavage fragments are indicated with an asterisk. The membrane was subsequently probed against *ACT2* and 18S rRNA as controls for the quality of the fractionation. **(B)** Impact of *EVD* over-proliferation in *epi15* with the *rdr6* as opposed to WT background. The total amount of seeds was counted in 7 to 8 individual plants from controls and homozygous WT or mutants from two individual F2 populations derived from crosses between *epi15* (carrying active EVD) and *rdr6* plants, as in Fig.6. Data points represent the median of two consecutive seed counts measurememnts for each individual plant. Table shows p-values (Holm-adjusted method) for pairwise comparison using Wilcoxon rank sum test (n.s: non significant*: p-value <0.05, **: p-value <0.01).

### Supplemental Table 1. Oligonucleotides used in this work.

**Genotyping primers**

| Target | Primer | Sequence 5'->3' | NOTES |
|---|---|---|---|
| ddm1-2 | ddm1 F | GCTGGAAGGGAAAGCTTAACAACCT | G-to-A mutation leads to the loss of a HaeIII restriction site: |
| (EMS mutant) | ddm1 R | acactgccatcgattctgcaaacc | wt: 80+121+218nt; ddm1: 121+298nt |
| dcl1-11 | dcl1-11 F | CGGGAATTTGTGAAGGAGGTTC | Use BIN-LB primer for mutant genotyping |
| (T-DNA insertion) | dcl1-11 R | CCTCTATCGCTCGTATTAACTC | (CGTCCGCAATGTGTTATTAAG) |
| hyl1-2 | hyl1-2 F | TTCTTGGAAATTGGATTGCAG | Use Salk T-DNA primer for mutant genotyping |
| (SALK_064863) | hyl1-2 R | AGTTCTCCCAGCGCTAATCTC | (ATTTTGCCGATTTCGGAAC) |
| xrn2-2 | xrn2 F | GCGCAAGTGGAGAAATCACT | Use Sail LB3 primer for mutant genotyping |
| (SAIL_781E02) | xrn2 R | TTTTGTCCCCATCTTCATGCC | (TAGCATCTGAATTTCATAACCAATCTCGATACAC) |
| xrn3-3 | xrn3 F | GCCTTCGATTTCAACAGGC | Use Sail LB3 primer for mutant genotyping |
| (SAIL_1172C07) | xrn3 R | GAAATCGAACACAAATCCG | (TAGCATCTGAATTTCATAACCAATCTCGATACAC) |
| xrn4-3 | xrn4 F | TCCCATGAGAGCCATGCATTC | Use Salk T-DNA primer for mutant genotyping |
| (SALK_020882) | xrn4 R | ACCATCCTGCAGGTCCAAGAA | (ATTTTGCCGATTTCGGAAC) |
| rdr6-12 | rdr6-12 F | CGTAATGAGCCTTGTTTTGG | 178 bp (-7bp for mutant allele) |
| (Fast neutron mutant) | rdr6-12 R | CGTCCTTGGGTTGGTTTCTTAG | |

**Northen blot probes**

| Target | Primer | Sequence 5'->3' |
|---|---|---|
| EVD GAG siRNA/mRNA | GAG F | TAAGTCAAGAAGACTTAGAGTTTA |
| | GAG R | ACTTTGCTCCTCATGAGTTTCTT |
| EVD "exon1" siRNA/mRNA | GAG-ex1 F | CGGCTAACAAAGGAGAAGTAGTGG |
| | GAG-ex1 R | CGGATTCTTTGAGTGAAGCAATGG |
| EVD "exon2" siRNA/mRNA | GAG-ex2 F | TATTGAGACGGGAAAAGTTATTGG |
| | GAG-ex2 R | CATCCGAATGAACTAGATCAAAAC |
| EVD intron siRNA/mRNA | GAG-in F | GTATCACTTTCTCATCACAAACATC |
| | GAG-in R | CTGAAAATACACATCATTAGGG |
| EVD RT (POL) siRNA | RT F | CAAAGACGGTATAGACTCTACCAAGAC |
| | RT R | CTCTATACTCCGATTCTGCACTCGAACA |
| EVD LTR siRNA | LTR F | TTGATCAAGACTCAAATAAGAAAGGCC |
| | LTR R | TATGCTCTGATACCATGAAAGAATAT |
| GFP siRNA/mRNA | GFP F | GCCGACAGTGGTCCCAAAGATG |
| | GFP R | TGATATCACTAGTGCGGCCG |
| GUS siRNA/mRNA | GUS F | GTGTGATATCTACCCGCTTCGCGTC |
| | GUS R | AAAGAGAGGTTAAAGCCGACAGCAGC |
| ACT2 mRNA | ACT2 F | GCACCCTGTTCTTCTTACCG |
| | ACT2 R | AACCCTCGTAGATTGGCACA |
| Athila6A siRNA | Athila6A F | caacgtcgatgaagctgaatcttgg |
| | Athila6A R | cctctcgaacttgggtttagttcc |
| Oligo probes | miR159 | TAGAGCTCCCTTCAATCCAAA |
| | miR171 | GATATTGGCGCGGCTCAATCA |
| | miR160 | TGGCATACAGGGAGCCAGGCA |
| | miR166 | GGGGAATGAAGCATGGTCCGA |
| | mir173 | GTGATTTCTCTCTGCAAGCGAA |
| | mir168 | TTCCCGACCTGCACCAAGCGA |
| | tasiR255 | TACGCTATGTTGGACTTAGAA |
| | U6 | AGGGGCCATGCTAATCTTCTC |

**5'OH RACE**

| Use | Primer | Sequence 5'->3' |
|---|---|---|
| 5' RNA adaptors | RNA adapter 5' dT | /5InvddT/ACGAUCAGUUCGCCGAUGCAG |
| | RNA adapter 3-P | ACCGCCGACGUCGAGGUGCCCGAAGGACCGCGCA-<br>-CCUGGUGCAUGACCCGCAAGCCCGGU/3Phos/ |
| EVD Gene-specific RT oligo | EVD_RT_Rv | CTAGCCTAGCATGCCACAAAG |
| Forward nested primers | Outer_Fw | ACCGCCGACGTCGAGGTGCC |
| | Inner_Fw | CGAAGGACCGCGCACCTGGT |
| EVD GAG region 1 RACE | EVD_R1_Out_Rv | AATCTTAGAACACACCTCATC |
| | EVD_R1_Inn_Rv | TTGGTAACTTCTCCGAC |
| EVD GAG region 2 RACE | EVD_R2_Out_Rv | CCGGCTTGACTTTGCTCCTC |
| | EVD_R2_Inn_Rv | GAGTTTCTTGAGAGAGATGAG |
| EVD GAG region 3 RACE | EVD_R3_Out_Rv | GACTTAGAGGAAAAACAGCTAG |
| | EVD_R3_Inn_Rv | AGAGTTTGGTGACAAGTCTTC |

**qPCR primers**

| Target | Primer | Sequence 5'->3' |
|---|---|---|
| Total GAG/EVD mRNA - | qGag-F | TTTGACCCGCGTGTTTGAAG |
| copy number | qGag-R | AATCTTCGGGTCAAGCGTTC |
| EVD flGAGPOL mRNA - | qRT-F | ACATATGGCTCCGACTACATGG |
| copy number | qRT-R | AGTTCGCCTTGGAGAAATGC |
| ACT2 mRNA | qACT2-F | GCACCCTGTTCTTCTTACCG |
| | qACT2-R | AACCCTCGTAGATTGGCACA |
| GAPC mRNA | qGAPC-F | ACTCAATCACTGCTACTCAG |
| | qGAPC-R | GTTGGGACACGGAAAGACATTCCA |
| RHIP1 mRNA | qRHIP1-F | GAGCTGAAGTGGCTTCAATGAC |
| | qRHIP1-R | GGTCCGACATACCCATGATCC |
| Spliced EVD/shGAG | qGAG spliced F | GTTGGTTGCTACATCCACACCT |
| mRNA | qGAG spliced R | TCAATATCCGGATTCTTTGAG |
| unspliced EVD/flGAG- | qGag unspliced F | GTTGGTTGCTACATCCACACCT |
| POL mRNA | qGag unspliced R | CAATTGAACTACTAGATGTTTGTGATG |
| Spliced GFP-EVDi- | qGFP spliced F | GCTGCTGGGATTACACATGGC |
| GUS/shGFP mRNA | qGFP spliced R | TCAATATCCGGATTCTTTGAG |
| Unspliced GFP-EVDi- | qGUS unspliced F | GCTGCTGGGATTACACATGGC |
| GUS/flGFP-GUS mRNA | qGUS unspliced R | CAATTGAACTACTAGATGTTTGTGATG |
| flGFP-GUS mRNA | qGUS F | GCACGGGAATATTTCGCGC |
| | qGUS R | GTATCGGTGTGAGCGTCGC |
| 18S rRNA | q18S F | TAGTTGGTGGAGCGATTTGTCTG |
| | q18S R | CTAAGCGGCATAGTCCCTCTAAG |
| U5 snoRNA | qU5 F | GAATACCGTGTGCTCTCCACGCT |
| | qU5 R | CCTCCAAAAATAGGCGTATGCCAC |
| MYB65 mRNA | qMYB65 F | GATGGTTCCTGATAGCCATACAGTTAC |
| | qMYB65 R | TAGGCATCAACAGAGTCAAGGAGATC |
| HAP2B mRNA | qHAP2B F | CTTGAACACTAAAAGTCAGAACTTGG |
| | qHAP2B R | GACACATTTAATCCGTTCGATAAGTT |
| TAS1c mRNA | qTAS1c F | TGTAGCGAAGAAGCATCA |
| | qTAS1c R | TGCAAAGCAAAACAGAAG |
| TAS3 mRNA | qTAS3 F | GAGACCGAAGTTTCTCCAAGGC |
| | qTAS3 R | CAGCACAGCGGATCCAATAATCTC |
| ARF4 mRNA | qARF4 F | ATACTACCCCACCCGGAAAC |
| | qARF4 R | TGAGACTGCATCGCAAAATC |
| HTT2 mRNA | qHTT2 F | GCCTGTCTAGCCTGTCTCGT |
| | qHTT2 R | CCCTCGGACTTATTCACTGC |
